# Biomarker-Aware Super-Resolution for 4D Flow MRI in a Synthetic Pulmonary-Artery Benchmark

**DOI:** 10.64898/2026.09.25.754495

**Authors:** Aitor Zubillaga-Unsain, Adrian Lluveras-Sires, Cesar Caballero-Gaudes, Jesús Ruiz-Cabello

## Abstract

**Purpose:** To evaluate, within a synthetic pulmonary-artery benchmark, whether biomarker-aware super-resolution improves hemodynamic endpoint fidelity in 4D flow MRI.

**Methods:** A 3D residual channel attention network (RCAN) was trained on 22 CFD simulations from 11 pulmonary-artery geometries with 2× spatial degradation. The original primary comparison evaluated full biomarker-aware training against a combined divergence–temporal-regularization control across 14 matched seeds on a fixed six-case test set. Additional matched analyses subsequently separated divergence and adjacent-frame temporal-difference regularization and evaluated biomarker supervision in a controlled 2×2 design with a common reconstruction schedule.

**Results:** In the original primary comparison, biomarker-aware training reduced PI error relative to the matched combined-regularization control (12.76 ± 2.34% vs. 17.12 ± 1.23%; paired difference −4.35 percentage points [95% CI −5.58, −3.09]; *p* = 0.0015). In the matched factorial analysis, biomarker supervision reduced PI error without (−2.235 percentage points [95% CI −3.514, −0.849]) and with (−4.125 percentage points [95% CI −5.392, −2.732]) combined regularization. PI improvement was not mirrored by uniformly improved reconstruction metrics: PSNR changed by −0.058 (*p* = 0.50) and +0.144 dB (*p* = 0.28), while SSIM decreased by 0.009 (*p* = 0.004) and 0.007 (*p* = 0.009), respectively. Divergence alone changed PI by +0.034 percentage points [95% CI −0.705, 0.809] while reducing DivRMS by 1.869 percentage points; temporal-difference regularization increased PI error by 1.562 percentage points [95% CI 0.594, 2.439].

**Conclusion:** Within this synthetic, fixed-case RCAN benchmark, biomarker supervision improved PI under both regularization conditions. PI improvement was not mirrored by uniformly improved reconstruction metrics. Temporal-difference—not divergence—regularization was the only isolated component associated with reproducible PI deterioration. These findings are configuration-specific and require geometry-independent and in vivo validation.

## 1 Introduction

Four-dimensional (4D) flow MRI is a phase-contrast technique that acquires three-directional velocity encoding time-resolved over the cardiac cycle, yielding volumetric velocity fields from which hemodynamic biomarkers can be derived.

Accurate hemodynamic biomarker quantification from 4D flow MRI — including peak systolic velocity, volumetric flow rate, and pulsatility index— is central to diagnosis and risk stratification in cardiovascular disease.^1,2,3,4^ In clinical practice, 4D flow acquisitions are typically limited to 2–3 mm isotropic voxels to maintain feasible scan times. This resolution introduces partial-volume averaging and can degrade peak-velocity and flow estimates in regions with sharp gradients.^5,6^

Deep-learning super-resolution (SR) offers a way to increase effective spatial resolution without prolonging acquisition time. Existing 4D flow SR methods and the broader SR literature have demonstrated strong reconstruction performance as measured by PSNR and SSIM.^7,8,9,10,11^ However, these metrics quantify reconstruction fidelity but do not directly measure the accuracy of clinical hemodynamic quantities.

Addressing this gap requires task-aligned objectives and physics-aware constraints. Biomarkers such as pulsatility index depend on localized velocity extrema, spatial integration over measurement planes, and temporal waveform structure. Small structured errors can materially affect clinical quantities despite acceptable voxel-level scores. Task-based evaluation frameworks have highlighted this issue in broader medical imaging contexts.^12^ Prior physics-informed work has improved flow plausibility, ^10,13^ but the relationship between loss design, physics regularization, and individual hemodynamic biomarker fidelity remains insufficiently characterized — particularly for pulsatility index, which is sensitive to both peak and mean flow estimates simultaneously. Because voxel-resolved hemodynamic ground truth is not available in vivo, controlled CFD-derived benchmarks remain useful for isolating whether loss design and validation-criterion choices affect endpoint fidelity under known flow conditions, consistent with prior PC-MRI/CFD validation work.^14^

Beyond loss design, a subtler issue is the choice of validation metric used for checkpoint selection during training. In image SR, checkpoint selection is typically driven by a single validation error such as PSNR; the analogous practice in velocity-field SR has often used peak-velocity error. When pulsatility index is the primary clinical target, selecting on a different metric can systematically favor checkpoints that are suboptimal for the actual endpoint of interest.

We evaluated, within a synthetic pulmonary-artery CFD benchmark used as a pulsatility-sensitive testbed, whether (1) explicit biomarker supervision improves pulsatility-index error relative to a matched combined divergence–temporal-regularization control and (2) changing the validation / early-stopping criterion from peak velocity to PI affects measured PI outcomes. The original four-arm analysis compared reconstruction-only, the historical combined-regularization control (combined divergence and adjacent-frame temporal-difference regularization), a biomarker-only control (biomarker supervision without divergence or temporal-difference regularization), and full biomarker-aware training across 14 matched seeds, with peak-velocity gradients disabled in the biomarker arms. Additional controlled analyses subsequently separated the two regularization components and tested biomarker supervision in a reconstruction-schedule-matched 2×2 design. Conventional reconstruction metrics were retained to characterize whether endpoint gains were accompanied by changes in voxel-level fidelity.

This work is a loss-design and evaluation-alignment study within a single RCAN backbone on a synthetic pulmonary-artery benchmark. It retains the original primary comparison of full biomarker-aware training against the matched combined-regularization control and adds controlled component and factorial analyses that isolate the effects of divergence, temporal-difference regularization, and biomarker supervision. Endpoint-aligned checkpoint reruns, a harder-noise screen, and a limited TemporalUNet4D probe further bound the findings. All conclusions are scoped to this benchmark; transferability beyond the evaluated configurations is not established.

## 2 Methods

### 2.1 Data Acquisition and Preprocessing

#### 2.1.1 Synthetic CFD Dataset

We used 22 synthetic pulmonary-artery bifurcation velocity fields generated by computational fluid dynamics (CFD). These are in-house numerical simulations of pulsatile incompressible Newtonian flow (*ρ* = 1060 kg/m^3^, *µ* = 3.5 × 10^−3^ Pa · s) in pulmonary-artery bifurcation geometries segmented from MRI; the CFD simulations were run in Ansys Fluent 2025 R2. The 22 simulations represented 11 underlying geometries with paired flow-regime realizations (11 laminar and 11 transitional). Each case provided a time-resolved three-component velocity field over a full cardiac cycle (20 phases). The executed clean primary and additional controlled configurations loaded isotropic 0.0020-m (2-mm) Cartesian reference volumes.

The present benchmark analysis used de-identified derived vessel geometries and synthetic CFD velocity fields; no identifiable participant images, clinical metadata, or intervention data are reported.

Mesh and solver settings: Each bifurcation geometry was meshed with a polyhedral core mesh and five prismatic boundary-layer inflation layers (first-cell height 0.02 mm, growth ratio 1.2; y^+^ < 1 for all cases). Final mesh sizes ranged from 1.2 to 2.1 million cells depending on vessel diameter. The pressure-based coupled solver was used with second-order spatial discretization and a bounded second-order implicit time scheme (time step = 5 × 10^−4^ s, 40 inner iterations per time step). Inlet boundary conditions were patient-representative pulsatile velocity profiles (parabolic with Womersley correction); outlet pressures were specified as zero gauge with a resistance boundary condition. Each case was run for three cardiac cycles; only the third cycle was retained to ensure periodic convergence. Velocity fields were exported at 20 evenly-spaced phases per cycle and resampled to the Cartesian reference volumes used for learning.

Each case included a vessel mask and cross-sectional planes at the inlet, a pre-bifurcation midline section, and the two branch outlets. Plane-based flow integration used a slab approximation over the vessel-mask intersection with physical voxel spacing.

The six held-out test cases are designated C1–C6 throughout the manuscript and supplementary material.

A stratified 12/4/6 train/validation/test split was fixed before any model comparison. The original split was performed at the simulation-case level rather than the geometry-group level; consequently, five geometry pairs were represented across different partitions. Accordingly, the test set should be interpreted as a held-out case set, not as a held-out-geometry set. All statistical comparisons use the same six test cases.

### 2.2 Low-Resolution Degradation Model

The controlled degradation mapped the 2-mm HR reference volumes to 4-mm LR inputs by 3D Gaussian point-spread-function blurring (σ factor 0.4) followed by 2× trilinear downsampling. The 4-mm LR input is coarser than the typical 2–3-mm clinical acquisitions used to motivate SR; this controlled 4-mm-to-2-mm task was not intended to reproduce the full clinical acquisition regime. Signal temporal averaging was applied over three consecutive frames at interior time points and over two frames at the temporal boundaries, according to the executed indexing. The additive noise itself was not temporally smoothed.

Gaussian noise was specified by

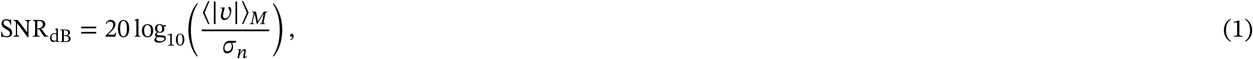

where ⟨|*v*|⟩_*M*_ is the vessel-mean velocity magnitude and σ_*n*_ is the noise standard deviation. SNR was sampled from 15– 30 dB during training and fixed per case at test time. Spatial correlation used a Gaussian kernel with σ = 0.5 LR voxels, reflect boundary handling, and global variance rescaling after filtering. All primary and additional controlled experiments used this same degradation; eight experiments compared noise-enabled and noise-free degradation with matched weight settings (Section 3.5). The degradation is therefore a controlled numerical benchmark rather than a simulation of the complete phase-contrast MRI acquisition chain.

### 2.3 Model Architecture

We employed a 3D RCAN ^8^ adapted for volumetric velocity-field reconstruction; temporal frames are processed independently with shared weights.

Architecture: Shallow feature extraction (3×3×3 conv, 64 channels), deep feature extraction via 5 residual groups × 10 Residual Channel Attention Blocks (RCAB; two 3×3×3 convs + channel attention with reduction ratio 16), 3D pixelshuffle upsampling (2×), and a global residual connection adding the trilinearly upsampled LR input to the output. Total parameters: 12.65M. For comparison, the 3D SRCNN baseline ^7^ has 0.41M parameters with trilinear pre-upsampling. Because integer 2× upsampling can differ by one voxel from the reference dimensions when an odd spatial dimension is present, RCAN outputs were resized to the reference target shape using nearest-neighbor interpolation whenever the pixel-shuffle output size did not match `target_size`. The reference HR tensor shape was supplied as `target_size` during training and evaluation, and this path was common to all compared RCAN arms. Retained diagnostics confirm activation in at least three test cases; complete prevalence across all 22 simulations could not be reconstructed because the frozen reproducibility archive did not retain spatial-shape metadata for every case.

Training hardware: NVIDIA A100-PCIe (80 GB VRAM) on the DIPC HPC hyperion cluster (Intel Xeon Platinum 8358 Icelake, 64 cores, 2 TB RAM). PyTorch 2.10.0 with CUDA 12.8.^15^ Mean wall-clock time: approximately 40 minutes per 250-epoch run (~10 s per epoch).

### 2.4 Loss Function Design

In all primary biomarker experiments, the peak-velocity term was inactive by design (*λ*_peak_ = 0); peak velocity was retained as a test-time clinical endpoint but contributed no training gradient.

The training objective combined seven loss terms with time-varying weights:

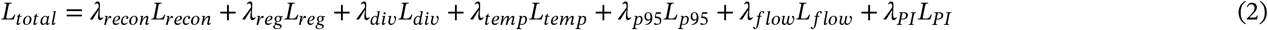

All seven *λ* values are specified in the progressive schedule (Table 1). Divergence regularization was active at a lower weight during the foundation phase and increased thereafter; temporal-difference and biomarker terms were introduced progressively in subsequent phases. Spatial Huber regularity was configured in the original arms but was held at zero in every additional component and factorial condition.

**Table 1.** Executed progressive loss-weight schedule for the full biomarker-aware arm. The values shown for interpolation rows vary linearly between the adjacent endpoints.

| Phase | Epochs | $\lambda_{\text{recon}}$ | $\lambda_{\text{reg}}$ | $\lambda_{\text{div}}$ | $\lambda_{\text{temp}}$ | $\lambda_{p95}$ | $\lambda_{\text{flow}}$ | $\lambda_{PI}$ |
| --- | --- | --- | --- | --- | --- | --- | --- | --- |
| Foundation | 1–41 | 1.0 | 0.1 | 0.1 | 0.0 | 0.0 | 0.0 | 0.0 |
| Interpolation | 42–51 | 1.0 → 0.8 | 0.1 | 0.1 → 0.3 | 0 → 0.2 | 0 → 0.1 | 0 | 0 |
| Physics | 52–141 | 0.8 | 0.1 | 0.3 | 0.2 | 0.1 | 0.0 | 0.0 |
| Interpolation | 142–151 | 0.8 → 0.5 | 0.1 | 0.3 → 0.25 | 0.2 → 0.25 | 0.1 → 0.2 | 0 → 0.2 | 0 → 0.2 |
| Final | 152–250 | 0.5 | 0.1 | 0.25 | 0.25 | 0.2 | 0.2 | 0.2 |

#### 2.4.1 Reconstruction Loss

Voxel-wise Charbonnier penalty ^16^ over masked voxels across all time frames:

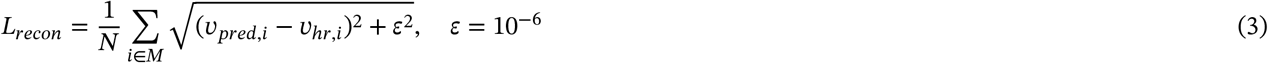

#### 2.4.2 Spatial Regularity Loss

Huber gradient penalty on predicted velocity field with spacing-aware finite differences (threshold *δ* = 0.1, applied per velocity component over all spatial axes):

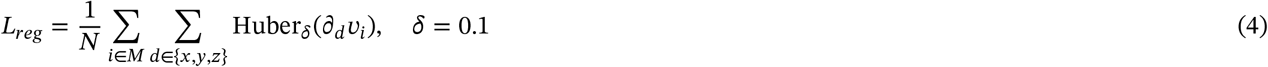

#### 2.4.3 Divergence and Temporal-Difference Regularization

Divergence loss: Dimensionless divergence penalty enforcing incompressibility (∇·v = 0), computed via second-order central finite differences with spacing-aware normalization:

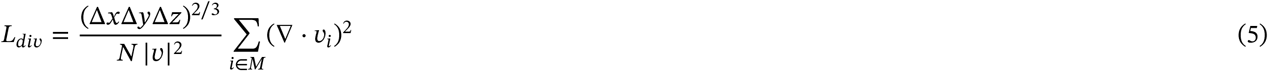

We use the geometric-mean voxel face area, (Δ*x*ΔyΔ*z*)^2/3^, to preserve dimensional consistency under voxel anisotropy. The normalized divergence is therefore dimensionally equivalent to a squared relative divergence per unit area. The mask is dilated by 1 voxel to include boundary voxels.

The adjacent-frame temporal-difference loss is a smoothness regularizer, not a conservation law. It is a first-order adjacent-frame penalty normalized by mean masked velocity magnitude:

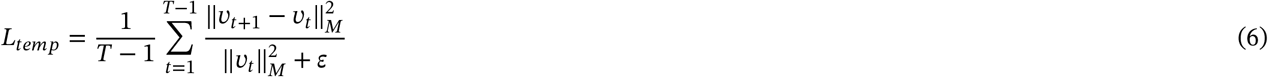

#### 2.4.4 Biomarker Loss

The biomarker loss directly optimizes flow rate and pulsatility index, and includes one active velocity-supervision term:

- Percentile-velocity term (*L*_*p*95_, final-phase *λ*_*p*95_ = 0.2): supervises the 95th-percentile velocity, providing a robust upper-distribution target.
- Peak-velocity term (*L*_peak_): the outer weight *λ*_peak_ is set to zero in all primary biomarker experiments. Peak velocity is evaluated at test time as a secondary clinical endpoint but contributes no gradient during training. This design choice is discussed in Section 4.4.

##### a. Percentile Velocity Loss

Percentile-based velocity supervision (95th percentile over the eroded mask) avoids the instability of optimizing the strict maximum:

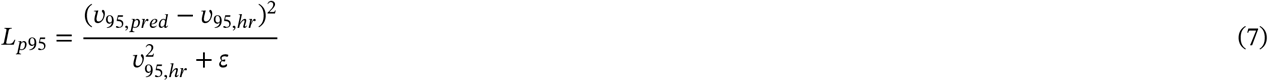

The peak-velocity loss (*L*_peak_, using the 99.5th percentile) follows the same formulation; in all primary biomarker experiments *λ*_peak_ = 0.

##### b. Flow Rate Loss

Volumetric flow rates computed at anatomically defined planes via slab integration:

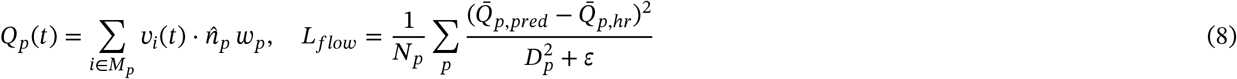

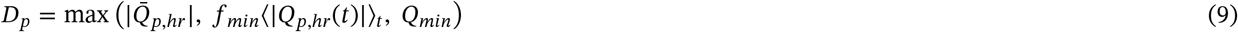

where *w*_*p*_ = (Δ*x*ΔyΔ*z*)/Δ*s*_min_ is the voxel face area (the product of the two spacings perpendicular to the plane normal 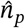), 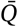 denotes the time-cycle mean, and *D*_*p*_ is a flow-rate scale with the same units as 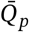, so *L*_*flow*_ is dimensionless. Hyperparameters: *f*_min_ = 0.05 and *Q*_min_ = 1 × 10^−6^ in the same flow-rate units as 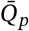; the stabilization constant ε is applied in the same squared units as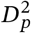.

##### c. Pulsatility Index Loss

The reported evaluation endpoint uses the conventional nonnegative definition *PI* = (*V*_*s*y*s*_ − *V*_*dia*_)/|*V*_*mean*_| per plane, where the three quantities are derived from the spatially averaged plane-normal velocity time curve. The executed training surrogate retained the orientation sign in the denominator, *PI*^*s*^ = (*V*_*s*y*s*_ − *V*_*dia*_)/*V*_*mean*_, and used

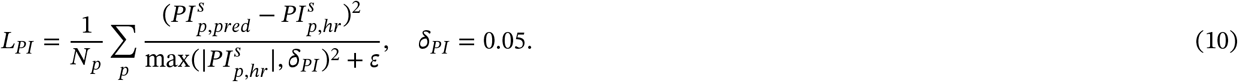

Plane normals had a consistent reference orientation for prediction and target. An end-to-end parity audit of the additional controlled analyses, comprising 2,016 unique run/case/plane observations, found no prediction–reference meanflow sign mismatches and a maximum conventional-versus-signed PI-error difference of 2.13 × 10^−6^ percentage points; no aggregate comparison, win/tie/loss count, confidence interval at reported precision, or statistical conclusion changed (Supplementary Note S14). The biomarker component of the final-phase total loss was

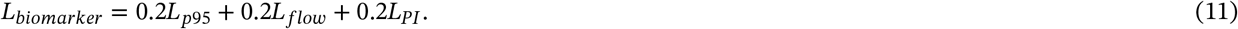

These coefficients are the final-phase *λ* values from the executed progressive schedule and are not normalized sub-weights within a separately scaled composite loss. The peak-velocity coefficient remained zero.

#### 2.4.5 Progressive Loss Scheduling

Training used a progressive three-phase schedule (Table 1).

The displayed boundaries reflect the actual scheduler indexing, including a one-epoch offset from the nominal configuration milestones. The reconstruction weight has a scheduled minimum of 0.5 and a hard floor of 0.1. In the original analysis, the reconstruction-only arm instead used fixed *λ*_*recon*_ = 1.0 with all other terms zero throughout 250 epochs. Because reconstruction scheduling differed across the original four arms, that comparison was not a clean 2×2 factorial design. The original primary full biomarker-aware versus combined-regularization comparison nevertheless used the same progressive reconstruction and regularization schedule in both arms, differing only in biomarker weights. The subsequent factorial experiment removed this broader four-arm imbalance by imposing a common schedule across all arms.

### 2.5 Training Protocol

#### 2.5.1 Optimization

We used the Adam optimizer ^17^ with an initial learning rate of 1×10^−4^, a 5-epoch linear warmup, and cosine annealing to 1 × 10^−6^ over 250 epochs. Gradient clipping (maximum norm 1.0) and mixed-precision training were applied throughout. Training used a batch size of 1, required because plane-based losses operate on full volumes, and validation was performed every 5 epochs.

#### 2.5.2 Early Stopping and Validation Criteria

Training used early stopping with patience of 40 epochs, retaining the best checkpoint according to a validation metric. Primary experiments and both additional controlled analyses used validation peak-velocity error as the checkpoint-selection criterion. The earlier matched-rerun analysis used the executed orientation-signed PI validation quantity; it was not a post hoc re-ranking of checkpoints from the original traces. Because those historical checkpoints were selected during training, no claim is made that counterfactual selection under a conventional-magnitude PI criterion would have chosen identical epochs.

No geometric augmentation was used; rotations and axis flips disrupt plane-referenced measurements. All experiments used fixed random seeds and requested deterministic PyTorch/cuDNN settings; because unsupported operations were permitted with warnings, bitwise identity across separate executions is not assumed. Per-run metadata included random seed, hardware, and software environment.

### 2.6 Evaluation Metrics and Baselines

#### 2.6.1 Primary Endpoints: Biomarker Errors

Evaluation used reference-geometry-derived masks and planes. Vessel masks came from the HR CFD/reference geometry, and cross-sectional planes were supplied from HR/Slicer geometry rather than inferred from the LR input or SR output. Plane origins and normals were transformed from LPS to RAS coordinates; normals were then normalized. Plane/vessel intersections were computed against the reference vessel mask using the physical voxel geometry. These masks and planes are evaluation aids and were not model inputs.

Mean absolute percentage errors (MAPE) were computed for the three hemodynamic endpoints:

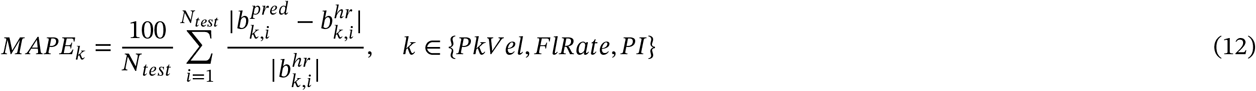

For PI, the conventional nonnegative value was calculated separately at each plane from the plane-normal velocity time curve. Absolute percentage error was calculated at each plane first, arithmetically averaged over the four planes to obtain a case-level PI error, and then arithmetically averaged across the six cases to obtain each run-level PI MAPE. The six cases were retained individually for descriptive summaries and were not pooled with seeds as independent observations. Reference PI values by case and plane are reported in Supplementary Table S13. Peak velocity used the 99.5th percentile over the eroded mask, aggregated as the mean of per-plane maxima.

Engineering reference thresholds were peak velocity < 5%, flow rate < 10%, and PI < 15%.

The 15% PI line is an engineering reference for the controlled benchmark, not a clinically validated pulmonary pass/fail threshold. Its supporting variability reference ^18^ derives from neurovascular rather than pulmonary 4D-flow MRI and is therefore contextual rather than anatomy-matched.

Secondary endpoints: divergence-RMS (normalized by mean velocity, %). Mass conservation error (signed inlet/outlet flow residual, %) was computed as a supplementary physics check; it is consistent with DivRMS trends and is not reported separately in the main results. Tertiary endpoints were PSNR, SSIM, and NMSE on velocity fields. The main tables prioritize PI and DivRMS; matched PSNR and SSIM contrasts are reported in Section 3.3, with the complete tertiary-metric results in Supplementary Table S10 and Supplementary Table S11. Gap closed (%) is defined as 100 × (ErrorLR − Errormodel)/ErrorLR; positive values indicate improvement over LR-native. Baselines: LR-native, trilinear interpolation, and 3D SRCNN, ^7^ all evaluated under identical conditions on the fixed 6-case test set.

#### 2.6.2 Statistical Analysis

##### Inferential unit and scope

The 14 matched seeds quantify stability to training stochasticity on the same fixed six-case test set and do not constitute 14 independent anatomical samples. Confidence intervals and tests therefore characterize seed-to-seed stability within this fixed benchmark, not patient-, anatomy-, or geometry-level generalization.

##### Original primary benchmark comparison

The primary comparison evaluates whether adding biomarker supervision improves PI relative to a matched combined divergence–temporal-regularization control across 14 seed-paired training runs on the fixed six-case test set. This is tested with a *two-sided* paired Wilcoxon signed-rank test on PI error differences (full biomarker-aware minus combined control); *n* = 14; *α* = 0.05.

##### Secondary comparison (contextual)

The full biomarker-aware vs. reconstruction-only comparison is presented as contextual evidence of the prior benchmark state, not as a second primary test. No formal *p*-values for secondary endpoints (peak velocity, flow rate, DivRMS) in the main-paper table; these are descriptive secondary endpoints.

##### Confidence intervals

95% CIs were computed by bootstrap resampling of the 14 paired seed differences (10,000 resamples, percentile interval, random seed 042). Matched-rerun confidence intervals in Table 5 used 1,000 bootstrap resamples, as stated in that table. These CIs describe stability across training instantiations on the fixed benchmark; they are not patient-level confidence intervals.

##### Per-case consistency

Per-case PI effects across the 6 test cases (mean ± SD of paired differences across 14 seeds) are reported in Supplementary Table S9 to characterize whether the primary improvement is consistent across cases or concentrated in 1–2 cases. These 6 cases represent within-benchmark case-level consistency, not generalization across independent patient samples.

##### Additional statistics

Effect size (Cohen’s *d*_*z*_ = mean/SD of paired differences; reported as the absolute magnitude for the primary comparison, and as signed values in the matched-rerun table where the sign indicates direction of improvement; rank-biserial *r* = 1 − 2*W*^+^/*W*_max_) and sign test (direction only, sensitivity analysis). All 14 seed pairs in the primary comparison were run under the same fixed protocol after the implementation correction; no sequential stopping rules were applied.

##### Integrity note (seed 047 tie)

In the primary benchmark comparison (full biomarker-aware vs. combined-regularization control), both seed-047 arms selected epoch 35 in the foundation phase, when all biomarker weights were zero and their active objectives were identical. Their archived validation value, test PI, and DivRMS were exactly equal; the structural tie (ΔPI = 0.0000 pp) is retained and counted as a tie.

##### Scope of secondary analyses

The harder-noise robustness screen (SNR [10,20] dB, 5 seeds, four arms) and TemporalUNet4D architecture-sensitivity probe (5 seeds, four arms, four-seed summary excluding seed 047 symmetrically) provide directional supporting evidence only. Neither constitutes a second primary study; no inferential *p*-values are reported for these analyses.

##### Additional controlled analyses

For the component experiment, seed-matched differences, percentile-bootstrap 95% CIs, two-sided paired Wilcoxon tests, and wins/ties/losses were calculated for RD−R, RT−R, RDT−R, RDT−RD, and RDT−RT. The divergence-by-temporal interaction was calculated per seed as (*PI*_*RDT*_ − *PI*_*RT*_) − (*PI*_*RD*_ − *PI*_*R*_). For the factorial experiment, the simple paired effects were 01−00, 11−10, 10−00, and 11−01; the biomarker-by-regularization interaction was (*PI*_11_ − *PI*_10_) − (*PI*_01_ − *PI*_00_). Interaction values were summarized by mean, SD, median, a 95% bootstrap CI, and a two-sided Wilcoxon signed-rank test. No seed×case pooling was used for inference.

### 2.7 Experiment Families

Eight experiment families were conducted with fixed seed sets and validation criteria. Families 1–6 comprise the original and supporting analyses. Families 7–8 are the subsequent controlled regularization-component and matched-factorial analyses; both retained the original data and evaluation pipeline while using a common progressive reconstruction schedule.

Table 2 summarizes the eight experiment families, seed sets, checkpoint criteria, and evidentiary roles.

**Table 2.**
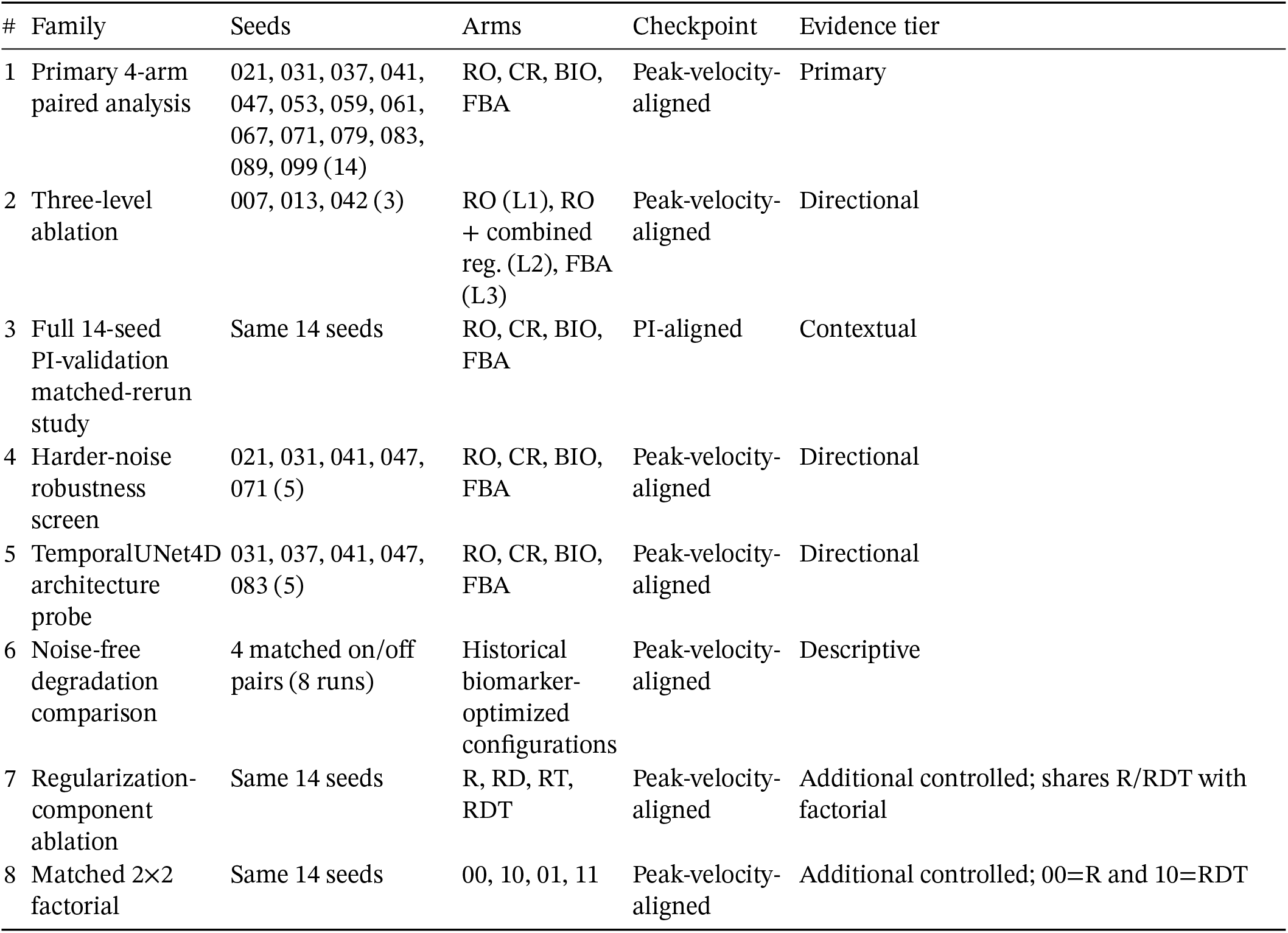
Experiment families. All RCAN families used 3D RCAN, 2× downsampling, no geometric augmentation, 250 epochs, batch size 1, and the fixed 12/4/6 case-level split. CR denotes the combined divergence–temporal-regularization control. Checkpoint criterion: peak-velocity-aligned = minimum validation peak-velocity error; PI-aligned = minimum executed orientation-signed validation PI error.

In the primary and post-correction supporting biomarker arms, the outer peak-velocity weight was set explicitly to *λ*_peak_ = 0; the historical Family 6 controls instead used nominally nonzero peak-velocity and other biomarker weights. Peak velocity was evaluated as a clinical endpoint at test time in both populations. The harder-noise screen used SNR [10,20] dB; all other families used SNR [15,30] dB. Supplementary Table S8 provides a compact provenance summary linking each experiment family to its evidence tier and reporting location. The corrected historical source audit is reported in Supplementary Table S7 and Supplementary Figure S1 and is retained for provenance only. The directional three-level ablation is shown in Supplementary Figure S2(a,b), whereas Supplementary Figure S3 provides the descriptive paired DivRMS visualization for the full biomarker-aware versus reconstruction-only comparisons.

For reference, the Supplementary Information comprises Supplementary Table S1, Supplementary Table S2, Supplementary Table S3, Supplementary Table S4, Supplementary Table S5, Supplementary Table S6, Supplementary Table S7, Supplementary Table S8, Supplementary Table S9, Supplementary Table S10, Supplementary Table S11, Supplementary Table S12, Supplementary Table S13, Supplementary Figure S1, Supplementary Figure S2(a,b), and Supplementary Figure S3; each is discussed in the corresponding Methods or Results section.

### 2.8 Four-Arm Paired Comparison (n = 14)

For each of 14 random seeds {021, 031, 037, 041, 047, 053, 059, 061, 067, 071, 079, 083, 089, 099}, four models were trained with the same split, architecture, degradation, and checkpoint criterion: reconstruction-only, combined-regularization control (CR), a biomarker-only control (biomarker supervision without divergence or temporal-difference regularization), and full biomarker-aware training. All runs used validation peak-velocity error for checkpoint selection. The combined-regularization control and full biomarker-aware arms shared the same progressive reconstruction and regularization schedule, so their primary pair differed only by biomarker supervision. Reconstruction scheduling was not identical across all four original arms, however, and the original four-arm display is therefore not interpreted as a factorial experiment. Archived run identifiers retain the historical physctl string for traceability.

The complete primary experiment matrix and per-seed four-arm results are reported in Supplementary Table S1 and Supplementary Table S2, respectively.

### 2.9 Additional Controlled Loss-Component Analyses

These analyses were performed after the original primary analysis to address the composition of the combined control and the reconstruction-schedule imbalance across the original four arms.

#### 2.9.1 Regularization-component ablation

Four matched RCAN conditions were evaluated: R (reconstruction only), RD (reconstruction + divergence), RT (reconstruction + adjacent-frame temporal-difference regularization), and RDT (reconstruction + both regularizers). Architecture, data and split, degradation, optimizer and learning-rate schedule, batch size, sampling, 250-epoch duration, common progressive reconstruction schedule, peak-velocity checkpoint criterion, evaluation pipeline, and all 14 seeds were identical. Spatial Huber regularity and all biomarker terms were zero. Divergence weights were 0.1, 0.3, and 0.25 in the foundation, physics, and final phases, respectively; temporal weights were 0, 0.2, and 0.25. No condition-specific tuning was performed.

#### 2.9.2 Matched 2×2 regularization-by-biomarker experiment

The factorial arms were 00 (combined regularization off, biomarker supervision off), 10 (combined regularization on, biomarker supervision off), 01 (combined regularization off, biomarker supervision on), and 11 (both on). Every arm used the same progressive reconstruction schedule, 250 epochs, optimizer and learning-rate schedule, architecture and initialization procedure, data order and degradation, batch size, validation frequency, peak-velocity checkpoint criterion, and 14 seeds. Spatial Huber regularity was zero in all four arms. When enabled, combined regularization used the component weights above. When enabled, biomarker weights were *λ*_*peak*_ = 0 throughout; *λ*_*p*95_ = 0/0.1/0.2, *λ*_*flow*_ = 0/0/0.2, and *λ*_*PI*_ = 0/0/0.2 across the foundation/physics/final phases. The executed schedule comprised foundation epochs 1–41, interpolation at 42–51, physics epochs 52–141, interpolation at 142–151, and the final phase from epoch 152. Factorial 00 reused the component R runs and factorial 10 reused the component RDT runs, with identical run, output, and checkpoint identities for every seed. The two analyses therefore share two arm sets and comprise six unique configurations across 14 seeds (84 trained models); RDT−R and 10−00 are the same underlying paired contrast under two analytical roles, not independent replication. Historical CR and FBA used spatial Huber regularity of 0.1, whereas it was intentionally zero in every factorial arm to isolate the two experimental factors. Because of this configuration difference, factorial 10 and 11 are not expected to reproduce the historical CR and FBA arm means.

### 2.10 Three-Level Ablation: Loss Configuration Effect on PI and DivRMS

A three-level ablation compared PI and DivRMS patterns across three training configurations:

1. Level 1 (reconstruction-only): Reconstruction + regularity only (seeds {007, 013, 042})
2. Level 2 (reconstruction + combined regularization): Level 1 + scheduled divergence + temporal consistency, biomarker terms = 0 (seeds {007, 013, 042})
3. Level 3 (full biomarker-aware): Full biomarker schedule with *λ*_*peak*_ = 0 (seeds {007, 013, 042})

Level 2 uses the same Foundation warm-start as Level 3 to avoid a training-protocol confound.

### 2.11 Matched-Rerun Analysis Using PI as the Validation Criterion (Full 14-Seed Study)

Matched reruns using PI as the validation / early-stopping criterion were completed for all four RCAN arms (reconstruction-only, combined-regularization control, biomarker-only control, and full biomarker-aware), each across all 14 primary seeds, enabling a direct within-seed comparison against the primary analysis (which used peak-velocity validation error). For the reconstruction-only and full biomarker-aware arms, five seeds were completed as an initial subset; the remaining nine seeds completed the full 14-seed set. For the combined-regularization control and biomarker-only control arms, the full 14-seed PI-aligned runs were conducted as a subsequent extension. All four PI-validation rerun sets used the same training configuration as their respective primary-arm runs, with only the validation / early-stopping criterion changed from peak velocity to pulsatility-index validation error.

### 2.12 Harder-Noise Robustness Screen

A five-seed (seeds 021, 031, 041, 047, 071) directional robustness screen evaluated all four arms under harder noise (SNR [10,20] dB instead of the primary [15,30] dB range). Results are descriptive (directional) and do not constitute a second primary study.

### 2.13 TemporalUNet4D Architecture-Sensitivity Probe

A five-seed (seeds 031, 037, 041, 047, 083) architecture-sensitivity probe used TemporalUNet4D (TUNet) instead of RCAN with the same four-arm loss design, training protocol, and data split. The TUNet backbone used three hierarchical levels with a base channel count of 32, temporal attention, and residual connections, at a 2× scale factor. Within each temporal block, a 3D spatial convolution was applied independently at each timestep, followed by a 1D convolution along the time axis. Seed 047 was unstable in the full biomarker-aware and biomarker-only control arms and is excluded symmetrically from all four arms in the four-seed summary (seeds 031, 037, 041, 083; *n* = 4). Seed 031 biomarker-only control is retained with a footnote noting elevated DivRMS and lower PSNR. This probe is a limited architecture-sensitivity check; it is not a head-to-head RCAN-vs-TUNet benchmark contest.

## 3 Results

Figure 1 summarizes the benchmark pipeline, four training arms, and the primary benchmark comparison used through-out the study.

**Figure 1.**
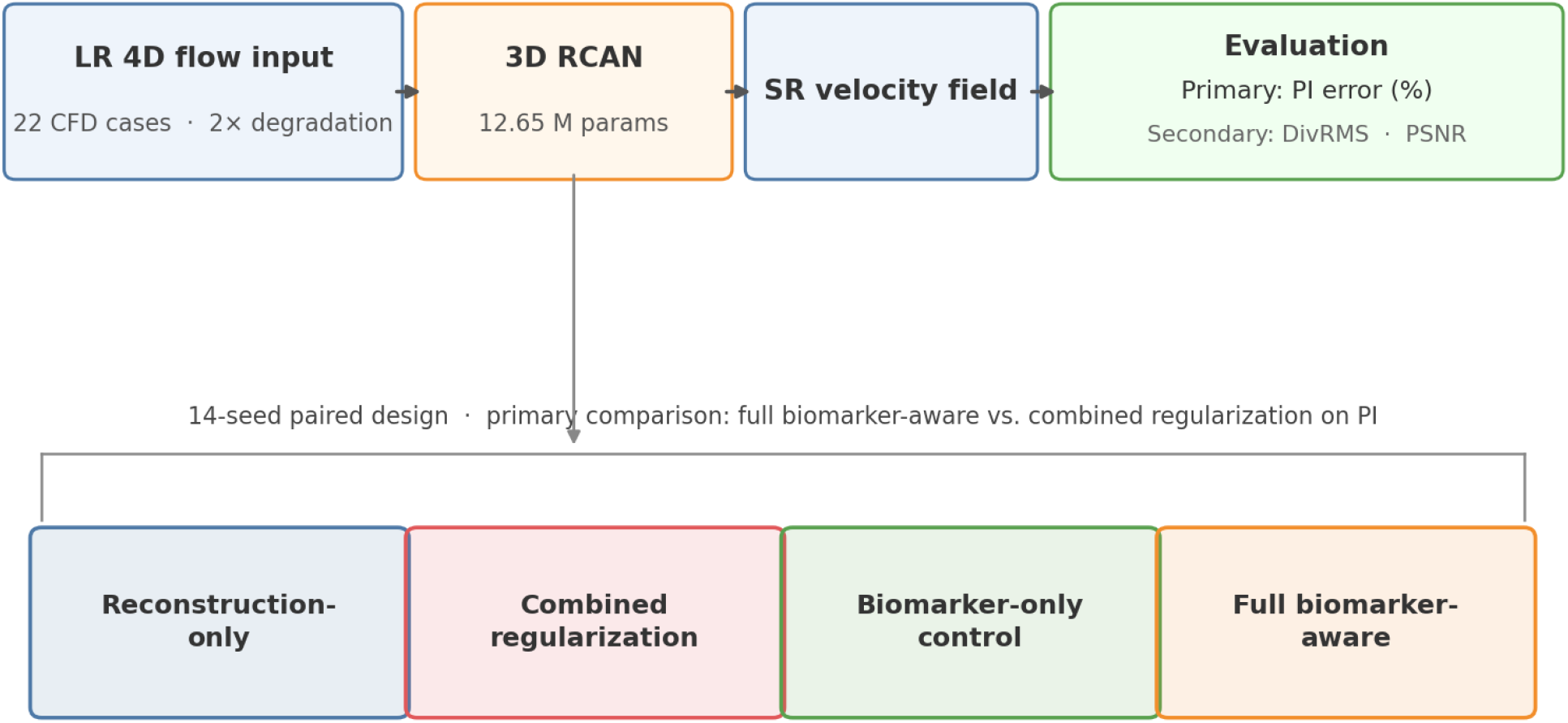
Original benchmark pipeline and four-arm training design. The 3D RCAN model reconstructs the HR velocity field from a 2× degraded input; evaluation uses the fixed six-case test set. The original primary comparison is full biomarker-aware versus the matched combined divergence–temporal-regularization control. The reconstruction-only arm used a different reconstruction schedule, so the historical four-arm display is not a factorial design; the subsequent matched factorial analysis is shown in Figure 4.

### 3.1 Four-Arm RCAN Results: PI and DivRMS Patterns Across Training Arms

Table 3 reports the four-arm results for the primary metrics. The PI row corresponds to the primary benchmark comparison (full biomarker-aware vs. combined-regularization control, two-sided Wilcoxon; all other rows are descriptive secondary endpoints). Figure 2(a) shows the PI results and Figure 2(b) shows the DivRMS results across the 14 matched seeds.

**Table 3.** Original four-arm RCAN paired analysis (*n* = 14 seeds). Values are mean MAPE (%) ± SD. **Bold PI row is the original primary comparison (full biomarker-aware vs. combined-regularization control, two-sided Wilcoxon** *p* = 0.0015). The combined-regularization arm contained divergence and temporal-difference regularization. This primary pair shared the same progressive reconstruction schedule; the four-arm set as a whole did not and is not interpreted factorially. Other rows are descriptive secondary endpoints.

| Metric | Recon.-only | Combined regularization | Biomarker-only control | Full bio-aware | Primary $\Delta$ (full bio-aware – combined regularization) [95% CI] |
| --- | --- | --- | --- | --- | --- |
| **PI (%)** | **14.22 ± 1.55** | **17.12 ± 1.23** | **13.12 ± 2.51** | **12.76 ± 2.34** | **−4.35 pp [−5.58, −3.09]; $p = 0.0015^\dagger$** |
| FlRate (%) | 5.10 ± 0.72 | 6.08 ± 1.12 | 5.46 ± 0.79 | 5.52 ± 0.66 | −0.56 pp (descriptive) |
| DivRMS (%) | 7.10 ± 0.14 | 5.31 ± 0.75 | 7.33 ± 0.14 | 4.94 ± 0.54 | −0.37 pp (descriptive) |
| PSNR (dB) | 39.99 ± 0.35 | 39.61 ± 0.39 | 39.75 ± 0.24 | 39.72 ± 0.22 | +0.11 dB (descriptive) |

<sup>†</sup> Two-sided paired Wilcoxon, $n = 14$ ; 13 wins / 1 tie / 0 losses. For seed 047, both arms selected epoch 35 in the foundation phase, where all biomarker weights were zero and the active objectives were identical. Their archived validation value, test PI (18.327%), and DivRMS (6.303%) were exactly equal; the single primary-comparison tie is therefore structural rather than a numerical near-miss. Seed 071 selected epoch 45 in both arms, during early interpolation when flow and PI supervision remained zero and percentile-velocity supervision had only begun to ramp; its outputs were near-identical but not equal, so no exact-tie mechanism is claimed. See Limitations.

**Figure 2.**
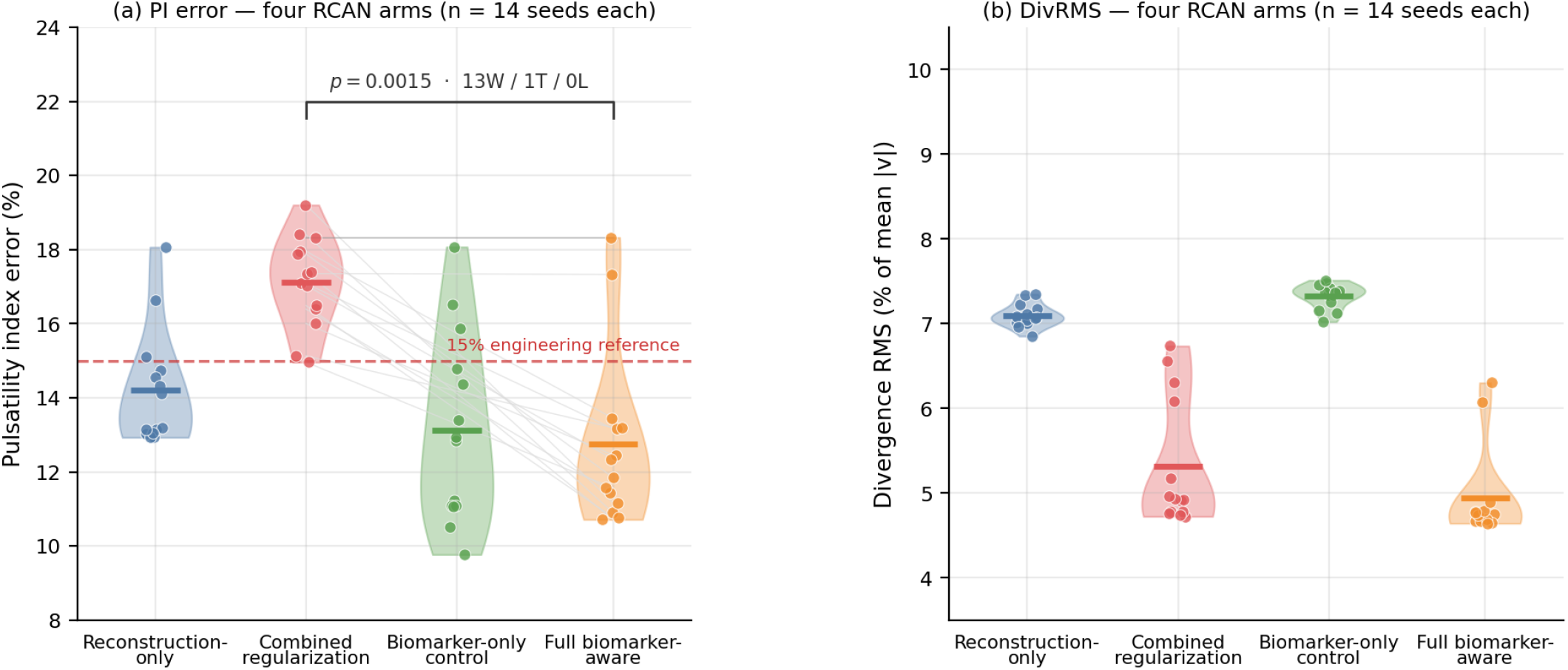
Original four-arm RCAN results (*n* = 14 seeds each). “Combined regularization” denotes combined divergence–temporal regularization, not conservation regularization alone. Connected lines show the original primary full-biomarker-aware versus combined-control comparison (*p* = 0.0015; 13 wins, 1 tie). The dashed 15% line is an engineering reference, not a clinically validated pulmonary threshold.

#### Primary benchmark result (full biomarker-aware vs. combined-regularization control)

Mean paired PI difference = −4.35 pp (SD = 2.47 pp), 95% bootstrap CI [−5.58, −3.09] pp (10,000 resamples, seed 042), two-sided Wilcoxon *p* = 0.0015, one-sided *p* = 0.0007 (sensitivity analysis only). Cohen’s *d*_*z*_ = 1.76 (magnitude of mean/SD of paired differences; large effect); rank-biserial *r* = 1.00.

#### Contextual secondary comparisons

1. *Full biomarker-aware vs. reconstruction-only:* Mean paired PI difference = −1.45 ± 1.39 pp, 95% bootstrap CI [−2.14, −0.75] pp; two-sided Wilcoxon *p* = 0.0052; *d*_*z*_ = 1.046 (magnitude); rank-biserial *r* = 0.810; 10/14 wins. The asymmetry between wins and losses is informative: the four losses are small (+0.05, +0.26, +0.51, +0.71 pp) while the ten wins range from −0.72 to −3.59 pp. This is contextual secondary evidence, not a co-primary test.
2. *Biomarker-only control vs. reconstruction-only:* Mean paired PI difference = −1.10 pp (SD = 1.48 pp), 95% bootstrap CI [−1.83, −0.35] pp; two-sided Wilcoxon *p* = 0.0203; 10/14 wins. Because reconstruction scheduling differed between these historical arms, this is contextual rather than factorial evidence.
3. *Combined-regularization control vs. reconstruction-only:* The historical combined divergence–temporal control had higher PI in all 14 seeds (mean +2.90 pp; *p* = 0.0001). This contrast neither isolates divergence from temporal smoothing nor controls the reconstruction-schedule difference; the subsequent matched experiments address both limitations.

Within the tested RCAN setting, the full biomarker-aware arm achieved the lowest mean PI error among the four arms. The DivRMS ordering (full biomarker-aware < combined-regularization control < reconstruction-only ≈ biomarker-only control) is consistent with an arm-level pattern in which combined regularization is most closely associated with divergence compliance in this benchmark; the biomarker-only control arm does not systematically improve DivRMS relative to reconstruction-only.

Relative to the 15% PI engineering reference, 12 of 14 full-biomarker-aware runs and 11 of 14 biomarker-only control and reconstruction-only runs were below the line, compared with 1 of 14 combined-control runs. This line is not a clinically validated pulmonary pass/fail threshold.

The PI results were also consistent across the fixed test cases: Figure 3(a) summarizes the conventional PI definition used for evaluation, while Figure 3(b) shows the descriptive per-case PI errors across the six cases.

**Figure 3.**
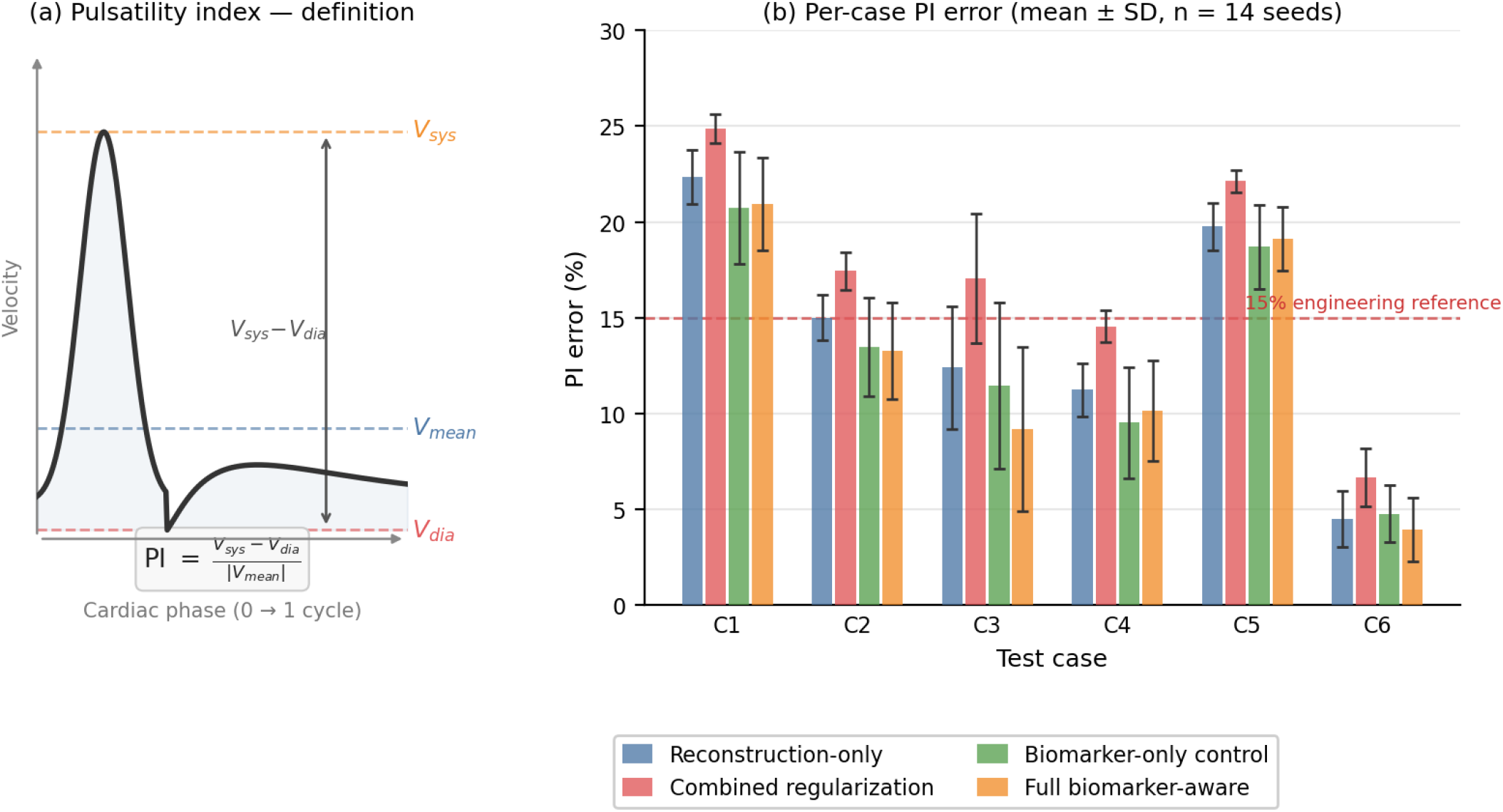
Conventional pulsatility-index definition and descriptive per-case consistency across the six fixed test cases. PI = (*V*_*s*y*s*_ − *V*_*dia*_)/|*V*_*mean*_|. Per-case errors are mean ± SD across seeds and are not treated as independent seed×case observations. The dashed 15% line is an engineering reference, not a clinically validated pulmonary threshold.

Reconstruction quality showed a small reduction in the full biomarker-aware arm: PSNR 39.72 ± 0.22 dB vs. 39.99 ± 0.35 dB for reconstruction-only (−0.27 dB); combined-regularization control PSNR = 39.61 ± 0.39 dB.

### 3.2 Regularization-Component Ablation

The matched component analysis localized the PI penalty of the original combined control (Figure 4(a,b)). Table 4 summarizes the arm-level PI and DivRMS results and the key paired PI differences from the additional controlled analyses. Divergence alone (RD vs. R) changed PI error by only +0.034 pp (95% CI [−0.705, 0.809]; *p* = 0.626), with 5 wins and 9 losses, while reducing DivRMS by 1.869 pp; all 14 seeds improved (*p* = 0.000122). Thus, divergence regularization substantially improved divergence compliance without reproducibly worsening PI when isolated.

**Table 4.**
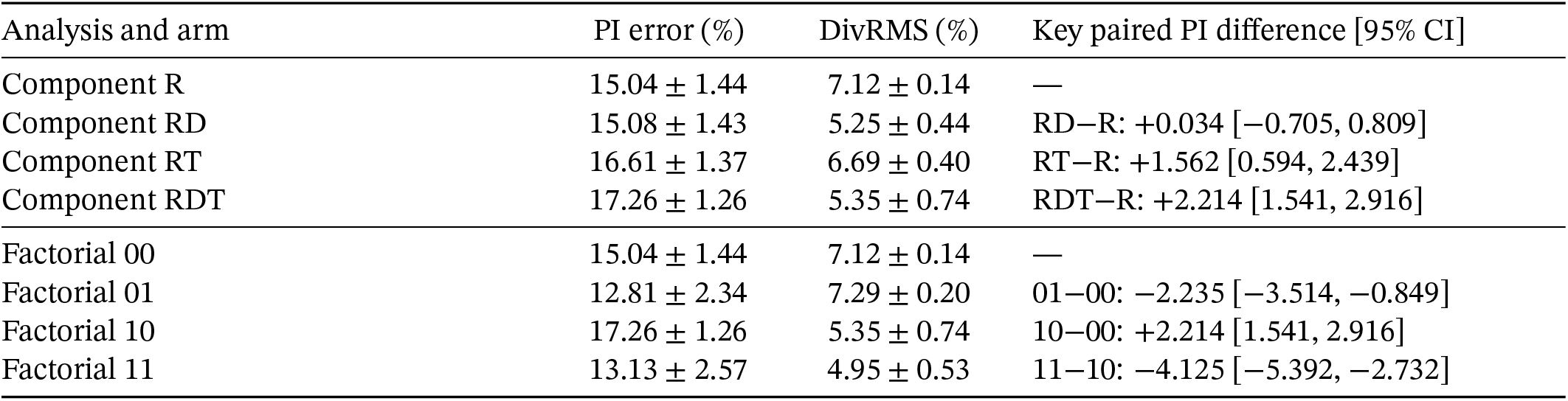
Additional matched controlled analyses (*n* = 14 seeds per arm). Values are mean ± SD. DivRMS is normalized by mean velocity magnitude and reported in percent. Factorial 00 reuses component R and factorial 10 reuses component RDT; RDT™R and 10™00 are therefore one underlying paired contrast shown in two analytical roles, not independent replications. Full secondary-metric and paired-comparison results are in Supplementary Table S10 and Supplementary Table S11.

**Figure 4.**
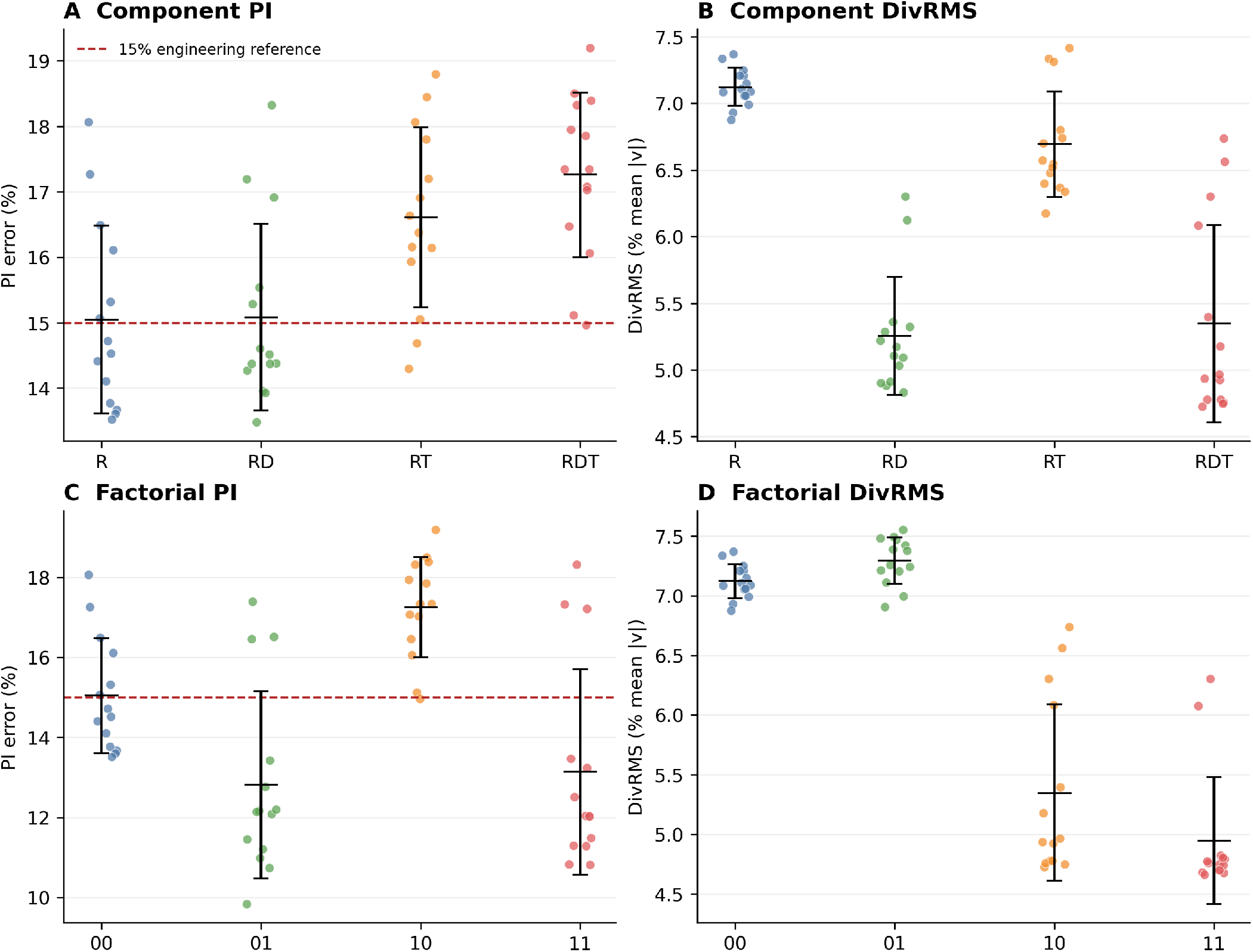
Additional matched controlled analyses across the same 14 training seeds and fixed six-case test set. **(a**,**b)** Regularization-component ablation for PI error and DivRMS. Divergence alone (RD) markedly reduced DivRMS without a reproducible PI change, whereas temporal-difference regularization (RT) increased PI error; the combined RDT condition had the largest PI penalty. **(c**,**d)** Matched factorial PI error and DivRMS for biomarker supervision off/on within combined-regularization strata. Biomarker supervision reduced PI in both strata, and combined regularization reduced DivRMS. Points denote seeds and bars denote mean ± SD. The dashed PI line is a 15% engineering reference, not a clinically validated pulmonary threshold.

Temporal-difference regularization alone (RT vs. R) increased PI error by 1.562 pp (95% CI [0.594, 2.439]; *p* = 0.013; 1 win, 1 tie, 12 losses). The combined RDT condition increased PI error by 2.214 pp relative to R (95% CI [1.541, 2.916]; *p* = 0.000122; all 14 seeds worsened) while reducing DivRMS by 1.774 pp in all 14 seeds. The formal divergence-by-temporal difference-in-differences was +0.619 pp (95% CI [−0.339, 1.789]; *p* = 0.345), providing no evidence of an interaction. Temporal-difference regularization was therefore the only isolated component associated with reproducible PI deterioration; the combined formulation produced the largest penalty, but its interaction was inconclusive.

### 3.3 Matched 2×2 Regularization-by-Biomarker Analysis

As shown in Figure 4(c,d), with the reconstruction schedule and all other optimization settings matched, biomarker supervision reduced PI error in both regularization strata. Without combined regularization, PI decreased from 15.04% (00) to 12.81% (01): paired difference −2.235 pp (95% CI [−3.514, −0.849]; *p* = 0.0067; 11/14 seeds improved). With combined regularization, PI decreased from 17.26% (10) to 13.13% (11): −4.125 pp (95% CI [−5.392, −2.732]; *p* = 0.0024; 12 wins, 1 tie, 1 loss).

Descriptive factorial PI results for each fixed test case are reported in Supplementary Table S12.

These PI gains were accompanied by small reproducible SSIM decreases (−0.009 for 01−00 and −0.007 for 11−10; 12/14 seeds unfavorable in each contrast). DivRMS increased modestly without combined regularization (01−00: +0.172 pp, 95% CI [0.105, 0.238]; *p* = 0.000366; 13/14 seeds unfavorable), whereas it decreased within the combined-regularization stratum (11−10: −0.402 pp, 95% CI [−0.771, −0.101]; *p* = 0.0131). These are bounded secondary tradeoffs and are not interpreted as an interaction.

Without biomarker supervision, combined regularization increased PI by 2.214 pp (10 vs. 00; 95% CI [1.541, 2.916]; *p* = 0.000122) while reducing DivRMS by 1.774 pp in every seed. With biomarker supervision, combined regularization changed PI by +0.325 pp (11 vs. 01; 95% CI [−1.261, 2.002]; *p* = 0.903) while reducing DivRMS by 2.348 pp in every seed (*p* = 0.000122). Peak-velocity error also decreased by 0.594 pp (*p* = 0.0419), a secondary finding.

The biomarker-by-regularization difference-in-differences was −1.889 pp (95% CI [−3.907, 0.154]; *p* = 0.0785). Thus, the estimated biomarker effect was numerically larger in the presence of combined regularization, but the formal interaction remained inconclusive. For seed 047, arms 10 and 11 both selected epoch 35, before biomarker activation, producing the expected exact tie; the seed was retained under the common checkpoint procedure.

In the post-correction matched factorial analysis, PI improvement was not mirrored by uniformly improved reconstruction metrics. For 01−00, PI improved by 2.235 pp while PSNR changed by −0.058 dB (*p* = 0.50) and SSIM by −0.009 (*p* = 0.004). For 11−10, PI improved by 4.125 pp while PSNR changed by +0.144 dB (*p* = 0.28) and SSIM by −0.007 (*p* = 0.009). The SSIM decreases were small in absolute terms but reproducible, with an unfavorable direction in 12 of 14 seeds in both contrasts (Supplementary Table S10 and Supplementary Table S11).

Descriptively, among the six unique controlled arm configurations, the highest mean test-set PSNR occurred in R, whereas the lowest mean PI error occurred in 01. This is an ordering observation across test-set means, not a model-selection analysis: checkpoints were selected by validation peak-velocity error, and cross-row comparisons are not paired inference. For descriptive scale context, Table 9 places LR-native at 34.2 dB and historical RCAN reconstruction-only at 39.99 dB, a difference of approximately 5.8 dB; by comparison, the six unique controlled-arm means in Supplementary Table S10 span 39.579–39.810 dB (0.23 dB).

### 3.4 Using PI as the Validation Criterion in Matched Reruns (Full 14-Seed Four-Arm Analysis)

A full 14-seed matched-rerun analysis using PI as the validation / early-stopping criterion was conducted across all four RCAN arms, enabling a direct within-seed comparison against the primary analysis (which used peak-velocity validation error). All four arms used the same training configuration as their respective primary-arm runs, with only the validation / early-stopping criterion changed to pulsatility-index validation error.

Full per-seed detail is reported in Supplementary Table S3, Panels A and B.

Figure 5 shows the per-seed ΔPI benefit of using PI rather than peak velocity as the validation / early-stopping criterion in matched reruns across all four RCAN arms. **Summary of results:** PI-validation reruns improved PI in all four arms across all or nearly all 14 seeds (Table 5):

1. *Full biomarker-aware arm:* Mean delta = −1.76 pp (95% CI [−3.00, −0.76] pp); two-sided Wilcoxon *p* = 0.0004; 13/14 seeds improved.
2. *Reconstruction-only arm:* Mean delta = −1.24 pp (95% CI [−2.07, −0.54] pp); two-sided Wilcoxon *p* = 0.0001; 14/14 seeds improved.
3. *Combined regularization arm:* Mean delta = −2.23 pp (95% CI [−3.01, −1.48] pp); two-sided Wilcoxon *p* < 0.0001; 14/14 seeds improved.
4. *Biomarker-only control arm:* Mean delta = −2.46 pp (95% CI [−3.67, −1.34] pp); two-sided Wilcoxon *p* = 0.0006; 13/14 seeds improved.

**Table 5.** Four-arm matched-rerun summary. Mean Δ = PI from matched reruns using PI as the validation / early-stopping criterion − PI from the corresponding primary runs using peak-velocity validation, averaged over 14 seeds (negative = PI-validation rerun better). 95% bootstrap CI (1000 resamples, percentile interval; the primary benchmark comparison used 10,000 resamples, Section 2.6). Cohen’s *d*_*z*_ is signed: negative values indicate the PI-validation rerun is superior. W/T/L = PI-validation rerun wins / ties / losses relative to the primary peak-velocity-validation run. All arms: *p* < 0.001 (two-sided Wilcoxon). Per-seed detail for all four arms in Supplementary Table S3.

| Arm | Mean $\Delta$ (pp) | 95% CI (pp) | W/T/L | $p$<br>(two-sided) | Cohen’s $d_z$ |
| --- | --- | --- | --- | --- | --- |
| Full biomarker-aware | $-1.76$ | $[-3.00, -0.76]$ | 13/0/1 | 0.0004 | $-0.80$ |
| Reconstruction-only | $-1.24$ | $[-2.07, -0.54]$ | 14/0/0 | 0.0001 | $-0.87$ |
| Combined regularization | $-2.23$ | $[-3.01, -1.48]$ | 14/0/0 | $<0.0001$ | $-1.49$ |
| Biomarker-only control | $-2.46$ | $[-3.67, -1.34]$ | 13/0/1 | 0.0006 | $-1.07$ |

**Figure 5.**
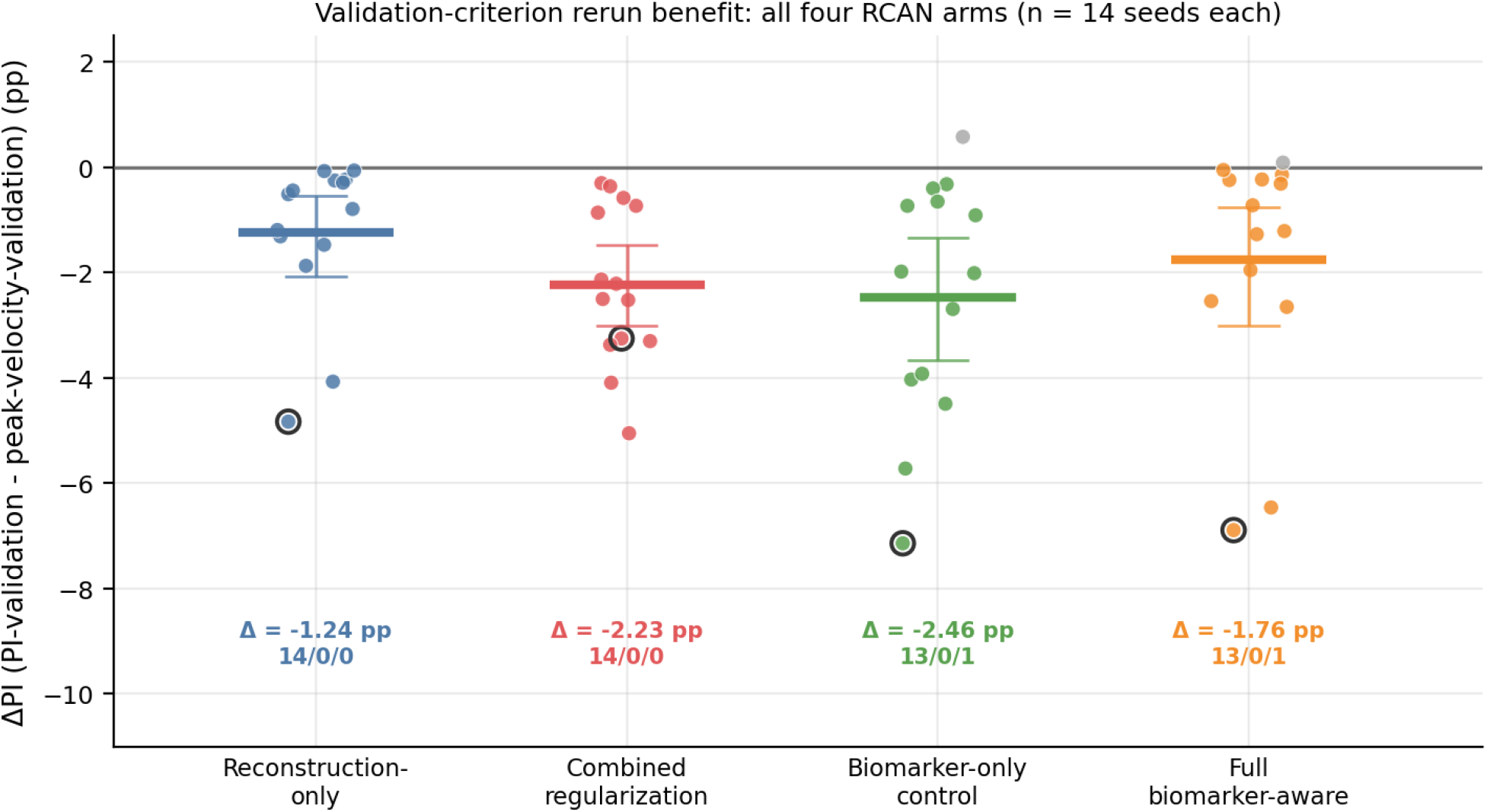
Validation-criterion rerun benefit: ΔPI = PI from matched reruns using PI validation − PI from the corresponding primary runs using peak-velocity validation, for all four RCAN arms (*n* = 14 seeds each). Negative Δ means the PI-validation rerun improved measured PI; positive Δ means it worsened PI. Each colored point is one seed; gray points are the few seeds where the PI-validation rerun did not help. Thick colored bars show the table-verified arm-level mean Δ (annotated beneath each arm with Δ and win / tie / loss count); vertical lines show the 95% bootstrap CI. The zero line marks no change. The bold-outlined point marks seed 047, the illustrative case where peak-velocity validation was most severely misaligned (largest improvement within the full biomarker-aware arm, Δ = −6.89 pp; also the largest improvement in the reconstruction-only and biomarker-only control arms). All four arms showed consistent improvement; full per-seed detail in Supplementary Table S3 and arm summary in Table 5.

Table 6 reports illustrative detail for five seeds (031, 037, 041, 047, 083) that were completed as the initial subset of the PI-validation matched-rerun study, including the case-study seed 047.

**Table 6.** Comparison of primary runs and matched PI-validation reruns for five illustrative seeds (full 14-seed per-arm analysis in Supplementary Table S3). “Peak-velocity-aligned checkpoint” = checkpoint selected in the primary run by minimum validation peak-velocity error; “PI-aligned checkpoint” = checkpoint selected in the matched rerun by minimum validation PI error. Δ (bio−recon) = full biomarker-aware PI minus reconstruction-only PI; negative values favor the full biomarker-aware model.

| Seed | FBA<br>peak-val.<br>PI (%) | $\Delta$ FBA–RO<br>peak-val.<br>(pp) | FBA<br>PI-val.<br>PI (%) | $\Delta$ FBA–RO<br>PI-val.<br>(pp) | Outcome |
| --- | --- | --- | --- | --- | --- |
| 031 | 11.45 | $-2.67$ | 11.31 | $-2.02$ | PI advantage retained |
| 037 | 10.90 | $-2.05$ | 10.66 | $-2.04$ | PI advantage retained |
| 041 | 11.16 | $-3.59$ | 10.93 | $-2.35$ | PI advantage retained |
| 047 | 18.33 | $+0.26$ | 11.44 | $-1.79$ | PI advantage gained |
| 083 | 11.85 | $-1.35$ | 11.54 | $-1.60$ | PI advantage retained, improved |

Seed 047 illustrates a validation-metric mismatch. In the primary run using peak-velocity validation, an early-epoch checkpoint was favored (low validation peak-velocity error but suboptimal PI), yielding test PI = 18.33% for the biomarker model vs. 18.06% for reconstruction-only, a marginal +0.26 pp loss. In the matched rerun using PI validation, the later checkpoint yielded test PI = 11.44% vs. 13.23% for reconstruction-only, a clear −1.79 pp improvement. Across all four RCAN arms, using PI as the validation / early-stopping criterion in matched reruns consistently improved measured PI, indicating that validation choice materially affects endpoint estimates in this benchmark regardless of which training loss was used.

### 3.5 Effect of Noise in the Degradation Model

In an auxiliary 8-experiment matched comparison, excluding noise from the degradation model substantially worsened peak-velocity error in the tested settings. Across the four matched standard-degradation pairs, the deterioration was 18.62–20.60 pp. The eight controls were historical pre-correction biomarker-optimized runs with a nominally nonzero but inoperative peak-velocity loss; within each noise-on/off pair, the training configuration and implementation were the same (Supplementary Table S6). This counterintuitive result indicates sensitivity of the learned pipeline to degradation and training conditions; it should not be interpreted as evidence that noise improves physical accuracy, and no causal mechanism is assigned without a dedicated experiment.

### 3.6 Harder-Noise Robustness Screen (SNR [10,20] dB, Directional)

A five-seed (seeds 021, 031, 041, 047, 071) four-arm screen evaluated all training arms under harder noise (SNR [10,20] dB). Table 7 reports mean ± SD across the five seeds for PI, DivRMS, and PSNR. Full per-seed values are reported in Supplementary Table S5.

**Table 7.** Harder-noise robustness screen (SNR [10,20] dB, *n* = 5 seeds). Values are mean ± SD across seeds. This is a directional screen only; no inferential statistics are reported. Bold PI ordering is directional.

| Arm | PI (%) | DivRMS (%) | PSNR (dB) |
| --- | --- | --- | --- |
| Reconstruction-only | $13.43 \pm 2.22$ | $7.97 \pm 0.21$ | $39.03 \pm 0.44$ |
| Combined regularization | $17.91 \pm 0.81$ | $6.30 \pm 0.48$ | $38.63 \pm 0.28$ |
| Biomarker-only control | **$10.18 \pm 1.05$** | $8.11 \pm 0.11$ | $38.96 \pm 0.07$ |
| Full biomarker-aware | $14.38 \pm 2.91$ | **$6.02 \pm 0.17$** | $38.65 \pm 0.16$ |

Across the five harder-noise seeds, the historical combined-control arm had higher PI and lower DivRMS than reconstruction-only in every case. This contrast does not isolate the regularization components. Biomarker-only control improved PI relative to reconstruction-only in all five seeds. The full biomarker-aware versus reconstruction-only comparison was mixed (3/5 wins; mean +0.95 pp favoring reconstruction-only). Thus, the primary PI benefit was not reproduced consistently in this limited harder-noise screen with the current hyperparameters.

### 3.7 Architectural Scope: Limited TemporalUNet4D Probe

Table 8 reports the TUNet four-arm results for four seeds (031, 037, 041, 083); seed 047 was excluded symmetrically from all four arms due to training instability in two arms (full biomarker-aware and biomarker-only control). Seed 031 (biomarker-only control arm) is retained but showed elevated DivRMS and lower PSNR relative to the other three seeds^∗^. Full five-seed detail is in Supplementary Table S4.

**Table 8.** TemporalUNet4D architecture probe: four-seed summary (seeds 031, 037, 041, 083; seed 047 excluded symmetrically due to instability in two arms). Values are mean ± SD. This four-seed summary is included to bound the architectural scope of the RCAN findings and should not be interpreted as a direct RCAN–TUNet comparison.

| Arm | PI (%) | DivRMS (%) | PSNR (dB) |
| --- | --- | --- | --- |
| Reconstruction-only | 12.30 ± 2.21 | 9.26 ± 0.45 | 37.37 ± 0.55 |
| Combined regularization | 18.56 ± 0.77 | 6.60 ± 0.08 | 37.72 ± 0.13 |
| Biomarker-only control* | 13.15 ± 0.43 | 10.80 ± 4.78 | 35.49 ± 2.04 |
| Full biomarker-aware | 18.18 ± 0.47 | 6.97 ± 0.50 | 37.66 ± 0.30 |

\*Seed 031 biomarker-only control contributed elevated DivRMS = 17.94% and lower PSNR = 32.46 dB, inflating the biomarker-only control SD.

Within TUNet, the combined-regularization and full biomarker-aware arms had lower DivRMS than reconstruction-only, but both had higher PI in the four-seed summary. The biomarker-only control arm most closely matched reconstruction-only PI. This ordering differs from the primary RCAN result and bounds its architectural scope; it does not isolate the effects of divergence and temporal terms in TUNet.

### 3.8 Descriptive Architecture and Baseline Context

This section places deterministic baselines and selected learned-model summaries on the same page for descriptive benchmark context only. Because the rows draw on heterogeneous evaluation bases, the table is not intended as a paired, inferential, or competitive method comparison.

Within this descriptive framing, the learned-model summaries show lower errors than the deterministic baselines on most reported endpoints, but these row-to-row differences should not be interpreted inferentially because the sample bases differ. SRCNN results are included only as a lightweight learned reference for the reconstruction-quality range. Using these descriptive summaries, RCAN full biomarker-aware closed approximately 58% of the peak-velocity gap, 55% of the flow-rate gap, and 24% of the PI gap relative to LR-native.

The gap-closed pattern differs qualitatively across endpoints. Trilinear interpolation alone recovered ~41% of the peak-velocity gap and ~40% of the flow-rate gap, but only ~5% of the PI gap. The descriptive endpoint summaries in Table 9 are the primary evidence for this observation.

**Table 9.** Descriptive architecture and baseline context within the fixed benchmark. *Deterministic baselines* (LR-native, trilinear): single evaluation on the fixed 6-case test set; no sampling variability. *Learned models*: SRCNN averaged over 5 seeds (lightweight learned reference included only to indicate a reconstruction-quality range); RCAN averaged over the primary 14-seed cohort. RO = reconstruction-only; FBA = full biomarker-aware. Seed-based rows report mean ± SD where shown; deterministic-baseline rows and SRCNN PSNR are reported as mean only. Pass = number of evaluations (or seed runs) meeting all three engineering targets simultaneously; denominators differ between rows. Because evaluation bases and sample sizes differ across all rows, no cross-row comparison should be interpreted as paired, inferential, or competitive.

| Model / sample | PkVel (%) | FlRate (%) | PI (%) | PSNR (dB) | Pass |
| --- | --- | --- | --- | --- | --- |
| *Deterministic baselines (single evaluation):* |  |  |  |  |  |
| LR-native (6 cases) | 13.2 | 12.4 | 16.8 | 34.2 | 0/1 |
| Trilinear (6 cases) | 7.8 | 7.4 | 15.9 | 37.1 | 0/1 |
| *Learned models (seed-based descriptive summaries only):* |  |  |  |  |  |
| SRCNN RO (5 seeds) | 5.5 ± 1.2 | 4.8 ± 0.9 | 14.8 ± 1.4 | 39.5 | 2/5 |
| RCAN RO (14 seeds) | 5.66 ± 1.49 | 5.10 ± 0.72 | 14.22 ± 1.55 | 39.99 ± 0.35 | 5/14 |
| RCAN FBA (14 seeds) | 5.56 ± 1.27 | 5.52 ± 0.66 | 12.76 ± 2.34 | 39.72 ± 0.22 | 5/14 |

## 4 Discussion

### 4.1 Endpoint Fidelity and Reconstruction Metrics

The matched controlled analyses provide the evidentiary basis for comparing endpoint and reconstruction behavior. Biomarker supervision improved PI in both factorial strata, but PSNR changed only slightly and inconsistently in direction, while SSIM decreased slightly in both contrasts. Thus, PI improvement was not mirrored by uniformly improved reconstruction metrics. This benchmark-specific result supports reporting endpoint errors alongside PSNR and SSIM.^12^

For peak velocity and flow rate, the gap-closed pattern in Table 9 shows that trilinear interpolation already recovered ~40% of the LR gap; RCAN reconstruction-only closed a further 16–19 pp. Pulsatility index shows a qualitatively different pattern: trilinear interpolation closed only 5% of the PI gap, and reconstruction-only SR closed ~15% total. The reason for this difference is an empirical observation in this benchmark; proposed explanations invoking PSF blur, waveform distortion, or temporal smoothing remain interpretive rather than demonstrated mechanistically.

### 4.2 Original Comparison and Controlled Loss-Component Analyses

In the original primary comparison (Table 3), full biomarker-aware training reduced PI error relative to the matched combined divergence–temporal-regularization control. The subsequent component ablation refined the interpretation of that control comparison. Divergence regularization alone substantially reduced DivRMS while leaving PI error essentially unchanged, whereas adjacent-frame temporal-difference regularization increased PI error. Thus, the deterioration in the original combined control should not be attributed to divergence regularization or “physics” generically. Temporal-difference regularization was the only isolated component associated with reproducible PI deterioration; the combined formulation produced the largest penalty, although the formal divergence-by-temporal interaction was inconclusive.

The clean 2×2 experiment also corrected the reconstruction-schedule imbalance across the original four arms. Biomarker supervision reduced PI error both without and with combined regularization. The direction of the clean 01−00 effect is consistent with the historical BIO-versus-RO contrast, while the historical comparison remains contextual because its reconstruction schedules differed. When biomarker supervision was present, adding combined regularization substantially improved DivRMS without a reproducible PI penalty. The biomarker effect was numerically larger in the combined-regularization stratum, but the formal interaction remained inconclusive. These results support biomarker supervision as a means of maintaining endpoint fidelity while incorporating regularization under this configuration; they do not establish synergistic interaction or independent replication of the shared RDT−R/10−00 contrast.

### 4.3 Flow-Rate Tradeoff and Peak-Velocity Accuracy

Mean flow-rate error remained below the 10% engineering target in all four arms (Table 3), ranging from 5.10% for reconstruction-only to 6.08% for combined-regularization control. Biomarker training introduced a slight average regression of 0.41 pp relative to reconstruction-only, calculated from the unrounded arm means (5.517% vs. 5.105%). This benchmark-specific pattern shows that PI optimization and mean-flow error did not move in parallel, but does not by itself identify the mechanism. In this setting, biomarker optimization improves the primary endpoint (PI) while introducing a modest average regression on a descriptive secondary endpoint (flow rate), with all arm means remaining below the engineering target.

Peak-velocity accuracy was similar across arms: mean full biomarker-aware 5.56% vs reconstruction-only 5.66%. Five of 14 seeds in each arm met the < 5% target. The fact that withholding peak-velocity gradient from training did not detectably worsen test-time peak-velocity accuracy is an informative secondary observation, though without formal equivalence testing we describe the difference only as “similar” or “not evidently harmed.”

### 4.4 Configuration Sensitivity of Biomarker Loss Optimization

In this synthetic pulmonary-artery benchmark, the PI improvement was observed under a specific set of training conditions. Each condition is associated with an empirical observation; the following explanations are interpretive rather than mechanistically demonstrated:

1. Zero peak-velocity gradient (*λ*_*peak*_ = 0): The percentile-velocity term *L*_*p*95_ (final-phase *λ*_*p*95_ = 0.2) supervises the upper velocity distribution without targeting the strict spatial maximum. This design may reduce competition between a single-voxel extremum target and the waveform-ratio objective, though this interpretation was not formally tested.

2. Foundation warm-start: An initial pre-biomarker phase preceded the introduction of biomarker gradients; under the executed scheduler, foundation weights applied through epoch 41, followed by a 10-epoch interpolation period. Empirically, skipping the warm-start created a reconstruction quality deficit in preliminary exploratory runs. This ordering is analogous to staged training strategies, but no formal ablation of the warm-start duration was performed in this benchmark.

### 4.5 Relationship Between PI and Divergence Compliance

The controlled analyses separate endpoint fidelity from divergence compliance more clearly than the historical arm labels. Divergence regularization consistently lowered DivRMS, including when isolated, but did not reproducibly alter PI on its own. Temporal smoothing increased PI error, plausibly because adjacent-frame penalties can attenuate pulsatile waveform variation; however, the present experiments establish an association for this specific formulation rather than a universal mechanism for all temporal regularizers. Biomarker supervision acted primarily on PI, while combined regularization retained its DivRMS benefit in the biomarker-enabled stratum. Image-level reconstruction metrics and endpoint errors therefore captured partly distinct behavior.

These arm-level patterns are specific to the primary RCAN conditions. The harder-noise screen and limited TUNet probe (Sections 3.6–3.7) did not reproduce the PI arm ordering consistently; the PI outcome associated with full biomarker-aware training should be treated as configuration-specific.

### 4.6 Validation Criterion and Matched-Rerun Implications

Seed 047 provides an illustrative example of validation-metric mismatch: in the primary run using peak-velocity validation, an early-epoch checkpoint was favored that was suboptimal for PI (see Table 6). The reconstruction-only arm result (14/14 seeds improved; Table 5) indicates that this mismatch with respect to PI arises in this benchmark under peak-velocity validation and is not specific to the biomarker training objective.

Within this benchmark, using PI rather than peak velocity as the validation / early-stopping criterion in matched reruns was better aligned with the stated primary endpoint. The combined-regularization control and biomarker-only control arms showed larger mean improvements than the full biomarker-aware and reconstruction-only arms (Table 5), suggesting that peak-velocity validation was more misaligned for the arms with highest absolute PI. Across all four RCAN arms, PI-validation reruns consistently reduced measured PI relative to the primary peak-velocity-validation runs, indicating that validation choice materially affects endpoint estimates in this benchmark regardless of the training loss used. If multiple endpoints are primary, a composite validation score weighted by endpoint priorities would be a reasonable extension.

### 4.7 Limitations and Future Work

#### Implementation transparency

An implementation defect in the peak-velocity loss function was identified in February 2026: the final aggregation step did not propagate gradients to the model parameters. Because all primary biomarker-arm configurations explicitly set *λ*_peak_ = 0, the affected term carried zero weight in the primary analysis; the reported primary results are therefore not expected to depend on this defect. All reported primary results were generated with the corrected implementation. Full technical details are provided in Supplementary Note S1.

#### Exploratory history and final analysis protocol

Approximately 119 exploratory runs were conducted before the implementation correction (February 2026), used for effect-size estimation and hyperparameter selection; they are not used as primary evidence (Supplementary Note S1). The 12/4/6 data split was fixed before any primary model comparison. Pre-analysis model-selection decisions were guided using validation-set performance, and test-set evaluation was reserved for the primary analysis after the implementation correction. All reported primary results are based exclusively on reruns performed after the implementation correction under a fixed protocol. An independently replicated analysis on a fully held-out dataset was not conducted.

#### Validation-set size

Hyperparameter and model-selection decisions rested on a validation partition containing four simulation cases, which limits the robustness of those selection decisions.

#### Early selected checkpoints (primary comparison)

For seed 047, both full biomarker-aware and combined-regularization arms selected epoch 35, within the foundation phase where all biomarker weights were zero and their active objectives and data order were identical. The archived validation value, test PI (18.327%), and DivRMS (6.303%) were exactly equal, as expected when both arms select a pre-activation checkpoint under the identical active objective; this is the single structural tie in the primary comparison, and it was retained without post-hoc exclusion. Seed 071 also selected epoch 45 in both arms, during early interpolation when flow and PI supervision remained zero and percentile-velocity supervision had only begun to ramp. Its outputs were near-identical but not equal, so no exact-tie mechanism is claimed.

#### Interaction estimates

Formal seed-wise difference-in-differences estimates were calculated in the additional controlled experiments. Neither the divergence-by-temporal interaction nor the biomarker-by-combined-regularization interaction excluded zero; claims of synergy are therefore not supported.

#### Scope — benchmark and architecture

This study evaluates a single RCAN backbone on a synthetic pulmonary-artery benchmark. Full architecture benchmarking was deferred. The SRCNN baseline provides a lightweight learned reference; it is not a state-of-the-art comparator. A limited five-seed TemporalUNet4D probe (Section 2.13; full per-seed data in Supplementary Table S4) found that the RCAN PI benefit was not reproduced under those conditions; it used a single hyperparameter configuration, symmetric exclusion of one unstable seed, and does not characterize the full TUNet configuration space or identify the cause of the PI ordering reversal.

#### Harder-noise screen limitations

The SNR [10,20] dB robustness screen used only five seeds and is directional evidence. The mixed result for full biomarker-aware vs. reconstruction-only under harder noise (3/5 seed wins, mean +0.95 pp favoring reconstruction-only) suggests that the PI benefit was not reproduced consistently in this harder-noise regime with the current hyperparameters.

#### Noise-free controls

The large deterioration in the single-seed noise-free controls was observed across all four matched PSF settings for peak-velocity error. Because removing additive noise also changes the degradation distribution presented during training, we interpret this result as evidence of pipeline sensitivity rather than evidence that noise is intrinsically beneficial; identifying the mechanism would require a dedicated degradation-domain ablation.

#### Temporal architecture

This study used RCAN with independent per-frame processing. A limited 5-seed TemporalUNet4D architecture-sensitivity probe was conducted (Section 2.13), but broader temporal-architecture benchmarking with systematic hyperparameter tuning remains deferred. Architectures with explicit temporal modeling may respond differently to the proposed loss design.

#### Synthetic data and acquisition realism

The executed clean experiments used 2-mm Cartesian CFD reference volumes with controlled degradation to 4-mm LR inputs. The LR input is coarser than typical 2–3-mm clinical acquisitions, while the 2-mm target lies at the fine end of that clinical range. The model did not simulate the complete phase-contrast MRI acquisition chain, including k-space sampling, VENC/velocity aliasing, background phase, eddy-current effects, or other scanner-and sequence-specific artifacts. The present degradation should therefore be interpreted as a controlled super-resolution benchmark rather than a realistic simulation of complete clinical 4D flow MRI acquisition.

#### Reference-assisted evaluation

Vessel masks and plane definitions were derived from HR CFD/reference geometry, and plane/vessel intersections used that geometry. These oracle evaluation aids were not inferred from LR or SR images. Performance when masks and planes must be estimated from acquired images was not assessed.

#### Geometry overlap across partitions

The fixed split was case-level rather than geometry-grouped: the 22 simulations represented 11 paired underlying geometries, and five geometry pairs crossed partition boundaries. Consequently, the present results do not establish generalization to previously unseen vascular geometries.

#### Test-set reuse

The same six-case test set was reused for follow-up analyses, including the component and factorial experiments. These analyses provide controlled mechanistic and sensitivity evidence but are not independent confirmatory evaluations on a previously untouched cohort.

#### Single anatomy

Experiments were restricted to pulmonary-artery bifurcations. Generalization to other anatomies requires multi-anatomy training and external validation.

#### Scale factor

Only 2× SR was evaluated. Higher factors may require different loss balancing.

#### Inferential unit and fixed test set

The experimental unit is the training seed on a shared six-case test set, not an independent anatomy or patient. Results describe consistency across training stochasticity and should not be interpreted as anatomical or patient-level inference.

#### Peak-velocity gradient

The primary analysis set *λ*_*peak*_ = 0 to avoid potential competition between the peak-velocity target and waveform-ratio objectives. Whether removing the percentile-velocity term *L*_*p*95_ as well would affect results was not tested.

#### Baseline headroom

Under extended training schedules (350 epochs), reconstruction-only models may already meet all engineering targets, leaving reduced margin for biomarker optimization to produce measurable improvement.

## 5 Conclusion

Within this synthetic pulmonary-artery RCAN benchmark, biomarker supervision reduced PI error in both the original primary comparison and a subsequently matched factorial experiment. In the matched factorial analysis, PI improvement was not mirrored by uniformly improved reconstruction metrics. Component ablation showed that temporal-difference regularization, rather than divergence regularization alone, accounted for most of the PI deterioration in the original combined control; divergence regularization improved divergence compliance without reproducibly worsening PI when isolated. These findings are specific to the evaluated synthetic, fixed-case benchmark and require validation on geometry-independent and in vivo data.

## Abbreviations

BIO: biomarker-only control (biomarker supervision without divergence or temporal-difference regularization)
CFD: computational fluid dynamics
CR: combined-regularization control
DivRMS: divergence root-mean-square
FBA: full biomarker-aware
HR: high-resolution
LR: low-resolution
MAPE: mean absolute percentage error
NMSE: normalized mean squared error
PI: pulsatility index
pp: percentage points
PSF: point spread function
PSNR: peak signal-to-noise ratio
RCAB: residual channel attention block
RCAN: residual channel attention network
RO: reconstruction-only
SNR: signal-to-noise ratio
SR: super-resolution
SSIM: structural similarity index
TUNet: TemporalUNet4D

## 6 Data Availability Statement

The data, code, and metadata that support the reported tables and figures are archived in a reproducibility package.^19^ The reproducibility package additionally includes the exact configurations, manifests, per-seed and per-case outputs, and aggregation scripts for the component and factorial analyses. Raw synthetic CFD velocity fields and trained model check-points are not publicly included because of source-data governance and storage constraints; they are available from the corresponding author upon reasonable request.

Supplementary material includes the original provenance and per-run tables plus full component and factorial metrics, paired comparisons, per-case PI summaries with reference PI values, scheduler/checkpoint notes, and the PI sign-parity audit.

## Ethics Approval and Consent to Participate

No new human participant data were collected for this benchmark study. Source MRI data used to derive the pulmonary-artery geometries were de-identified before CFD simulation; the present analysis used only de-identified derived vessel geometries and synthetic CFD velocity fields. No identifiable participant images, clinical metadata, or intervention data are included in the manuscript or supplementary material. Consent for publication is not applicable because no identifiable participant information is reported.

## 7 Acknowledgments

This work was supported by ELKARTEK 2023 (CIC biomaGUNE and BCBL) and by the Basque Government, Department of Science, Universities and Innovation, through the IKUR 2030 Strategy, under the IKUR-HPC&AI 2025–2026 project SPADE-MRI (Synthetic Data Pipeline for Accelerated 4D Flow MRI: Enabling Robust Deep Learning Models). Jesús Ruiz-Cabello is funded by grant PID2024-155807OB-C21, funded by MCIN/AEI/10.13039/501100011033, by “ERDF A way of making Europe,” and by the European Union; grants DTS24/00043 and ISCIII-AES-2024/000225, funded by Instituto de Salud Carlos III (ISCIII); ELKARTEK 2025 (KK-2025-00084); and the Fundación contra la Hipertensión Pulmonar. This work acknowledges the use of ICTS-ReDIB, supported by the Ministry of Science, Innovation and Universities (MICIU) at BioImaC and CIC biomaGUNE. Computational resources were provided by the DIPC High-Performance Computing facility. We thank the DIPC HPC support team and collaborators at CIC biomaGUNE for technical discussions related to 4D flow MRI biomarker reproducibility.

The authors used OpenAI Codex/ChatGPT as an editorial and LATEX assistance tool during manuscript preparation, including language polishing, table-formatting checks, and graphical-abstract layout support. All AI-assisted outputs were reviewed and edited by the authors, who take full responsibility for the manuscript content.

## Author Contributions

Aitor Zubillaga-Unsain: Conceptualization, Methodology, Software, Investigation, Formal analysis, Data curation, Visualization, Writing–original draft, Writing–review & editing. Adrian Lluveras-Sires: Writing–review & editing. Cesar Caballero-Gaudes: Funding acquisition, Resources, Supervision, Writing–review & editing. Jesús Ruiz-Cabello: Funding acquisition, Resources, Supervision, Writing–review & editing.

## References

1. Markl M, Frydrychowicz A, Kozerke S, Hope M, Wieben O. 4D flow MRI. J Magn Reson Imaging. 2012;36(5):1015–36. doi: 10.1002/jmri.23632.

2. Dyverfeldt P, Bissell M, Barker AJ, Bolger A, Carlhall CJ, Ebbers T, et al. 4D flow cardiovascular magnetic resonance consensus statement. J Cardiovasc Magn Reson. 2015;17(1):72. doi: 10.1186/s12968-015-0174-5.

3. Barker AJ, Lanning C, Shandas R. Quantification of hemodynamic wall shear stress in patients with bicuspid aortic valve using phase-contrast MRI. Ann Biomed Eng. 2010;38(3):788–800. doi: 10.1007/s10439-009-9854-3.

4. Bissell MM, Raimondi F, Ait Ali L, et al. 4D Flow cardiovascular magnetic resonance consensus statement: 2023 update. J Cardiovasc Magn Reson. 2023;25(1):40. doi: 10.1186/s12968-023-00942-z.

5. Pravdivtseva MS, Gaidzik F, Berg P, Ulloa P, Larsen N, Jansen O, et al. Influence of Spatial Resolution and Compressed SENSE Acceleration Factor on Flow Quantification with 4D Flow MRI at 3 Tesla. Tomography. 2022;8(1):457–78. doi: 10.3390/tomography8010038.

6. Cherry M, Khatir Z, Khan A, Bissell M. The impact of 4D-Flow MRI spatial resolution on patient-specific CFD simulations of the thoracic aorta. Sci Rep. 2022;12(1):15128. doi: 10.1038/s41598-022-19347-6.

7. Dong C, Loy CC, He K, Tang X. Image super-resolution using deep convolutional networks. IEEE Trans Pattern Anal Mach Intell. 2016;38(2):295–307. doi: 10.1109/TPAMI.2015.2439281.

8. Zhang Y, Li K, Li K, Wang L, Zhong B, Fu Y. Image super-resolution using very deep residual channel attention networks. In: Computer Vision – ECCV 2018; 2018. p. 294–310. doi: 10.1007/978-3-030-01234-2_18.

9. Ferdian E, Suinesiaputra A, Dubowitz DJ, et al. 4DFlowNet: Super-Resolution 4D Flow MRI Using Deep Learning and Computational Fluid Dynamics. Front Phys. 2020;8:138. doi: 10.3389/fphy.2020.00138.

10. Fathi MF, Perez-Raya I, Baghaie A, et al. Super-resolution and denoising of 4D-Flow MRI using physics-informed deep neural nets. Comput Methods Programs Biomed. 2020;197:105729. doi: 10.1016/j.cmpb.2020.105729.

11. Shit S, Zimmermann J, Ezhov I, Paetzold J, Sekuboyina A, Kofler F, et al. SRflow: Deep learning based super-resolution of 4D-flow MRI data. Front Artif Intell. 2022;5:928181. doi: 10.3389/frai.2022.928181.

12. Barrett HH, Abbey CK, Clarkson E. Objective assessment of image quality. III. ROC metrics, ideal observers, and likelihood-generating functions. J Opt Soc Am A Opt Image Sci Vis. 1998;15(6):1520–35. doi: 10.1364/JOSAA.15.001520.

13. Raissi M, Perdikaris P, Karniadakis GE. Physics-informed neural networks: A deep learning framework for solving forward and inverse problems involving nonlinear partial differential equations. J Comput Phys. 2019;378:686–707. doi: 10.1016/j.jcp.2018.10.045.

14. Puiseux T, Sewonu A, Meyrignac O, Rousseau H, Nicoud F, Mendez S, et al. Reconciling PC-MRI and CFD: An in-vitro study. NMR Biomed. 2019;32(5):e4063. doi: 10.1002/nbm.4063.

15. Paszke A, Gross S, Massa F, et al. PyTorch: An Imperative Style, High-Performance Deep Learning Library. In: Advances in Neural Information Processing Systems (NeurIPS); 2019. doi: 10.48550/arXiv.1912.01703.

16. Charbonnier P, Blanc-Féraud L, Aubert G, Barlaud M. Two deterministic half-quadratic regularization algorithms for computed imaging. Proceedings of 1st International Conference on Image Processing. 1994;2:168–72. doi: 10.1109/ICIP.1994.413553.

17. Kingma DP, Ba J. Adam: A Method for Stochastic Optimization. In: International Conference on Learning Representations (ICLR); 2015. doi: 10.48550/arXiv.1412.6980.

18. Rivera-Rivera LA, Roberts GS, Peret A, et al. Unraveling diurnal and technical variability in cerebral hemodynamics from neurovascular 4D-Flow MRI. J Cereb Blood Flow Metab. 2024;44(8):1362–75. doi: 10.1177/0271678X241232190.

19. Zubillaga-Unsain A, Lluveras-Sires A, Caballero-Gaudes C, Ruiz-Cabello J. Reproducibility package for Biomarker-Aware Super-Resolution for 4D Flow MRI in a Synthetic Pulmonary-Artery Benchmark. Zenodo. 2026. Available from: https://doi.org/10.5281/zenodo.22099891. doi: 10.5281/zenodo.22099891.

