## Supplementary Material for "Biomarker-Aware Super-Resolution for 4D Flow MRI in a Synthetic Pulmonary-Artery Benchmark"

Preprint. This manuscript is under peer review.

#### Contents

**Supplementary Note S1: Peak-Velocity Gradient-Flow Defect and Fix** 2

**Supplementary Table S1: Primary 14-Pair Experiment Matrix** 4

**Supplementary Table S2: Per-Seed Four-Arm Results** 5

**Supplementary Table S3: Validation-Criterion Rerun Comparison** 7

**Supplementary Table S4: TemporalUNet4D Probe** 9

**Supplementary Table S5: Harder-Noise Robustness Screen** 11

**Supplementary Table S6: Noise-Enabled vs. Noise-Free Degradation Controls** 12

**Supplementary Table S7: Historical Exploratory PSNR–PI Source Data** 13

**Supplementary Table S8: Experiment-Family Provenance Summary** 15

**Supplementary Table S9: Per-Case PI Consistency** 16

**Supplementary Table S10: Full Arm Summaries** 17

**Supplementary Table S11: Full Paired Comparisons** 18

**Supplementary Table S12: Fixed-Test-Case PI Results** 20

**Supplementary Table S13: Reference PI Values** 21

**Supplementary Note S14: Scheduler, Checkpoint, and PI Sign-Parity Details** 22

**Supplementary Figure S1: Historical Exploratory PSNR–PI Scatter** 24

**Supplementary Figure S2: Three-Level Ablation** 25

**Supplementary Figure S3: Paired DivRMS Scatter** 26

### Supplementary Note S1: Peak-Velocity Gradient-Flow Defect and Fix

**Summary for main text (also stated in §4.7):** An implementation defect in the peak-velocity loss function was discovered in February 2026. Because all primary biomarker-arm configurations explicitly set $\lambda_{\text{peak}} = 0$, the affected term carried zero weight in the primary analysis; the reported primary results are therefore not expected to depend on this defect. All reported primary results were produced with the corrected implementation. This note summarizes the technical validation.

#### S1.1 Defect Description

A retrospective review identified a defect in the peak-velocity loss: intermediate velocity values were processed in a way that interrupted gradient propagation from this loss term back to the model parameters. As a result, the final aggregation step for peak velocity no longer transmitted gradient, and the backward pass contributed zero gradient through that component.

**Affected component:** The peak-velocity loss function. The defect arose during the velocity-reduction and final aggregation steps, where a differentiable path to the model parameters was no longer preserved.

**Unaffected components:** The percentile-velocity, flow-rate, and pulsatility-index loss functions preserved differentiable aggregation throughout and were not affected.

#### S1.2 Fix

The peak-velocity loss was revised so that all reduction and aggregation steps preserve a differentiable path from the loss term to the model parameters, restoring gradient propagation through that component.

#### S1.3 Validation

Validation comprised 16 dedicated checks of gradient propagation spanning all four biomarker loss functions, reduction methods, plane scopes, normalization modes, and gradient magnitudes. All 109 implementation checks passed.

#### S1.4 Impact on Reported Results

**Why the primary analysis is clean (two independent reasons):**

1. **Design-level protection:** All primary biomarker arm configurations explicitly set $\lambda_{\text{peak}} = 0$. Peak-velocity loss therefore contributed no gradient by design, not because of the defect.
2. **Execution after correction:** The Primary 14-pair four-arm analysis, Directional three-level ablation, and Contextual validation-criterion rerun studies were all run with the corrected implementation.

**Exploratory runs before the implementation correction** (approximately 119 runs with biomarker loss, conducted before 2026-02-17) are not used as primary evidence anywhere in the main text. These exploratory runs informed effect-size estimation and study design. The eight seed-042 degradation controls are additionally retained as descriptive historical sensitivity evidence in Supplementary Table S6. Supplementary Table S7 and Figure S1 are retained solely as an audited record of a historical exploratory screen and do not support the conclusions.

##### S1.5 Experiment Sequence

After the defect was identified, corrected, and validated, the primary analysis was rerun under the corrected fixed protocol. The post-correction primary and supporting program comprised reruns of the full biomarker-aware and reconstruction-only arms of the primary 14-seed comparison (28 runs); the nine-run three-level ablation; an initial five-seed set of PI-validation reruns for those two arms (10 runs); reruns of the combined-regularization control and biomarker-only control arms (14 runs each); PI-validation reruns for the remaining nine reconstruction-only and full biomarker-aware seeds (18 runs); the harder-noise robustness screen (20 runs); the TemporalUNet4D architecture probe (20 runs); and PI-validation reruns for the combined-regularization control and biomarker-only control arms (14 runs each). The eight seed-042 noise-enabled/noise-free control runs in Supplementary Table S6 were executed on 2026-02-09 through 2026-02-12, before the 2026-02-17 implementation correction. The later controlled component and factorial analyses used 84 unique trained models: factorial 00 reused component R and factorial 10 reused component RDT for all 14 seeds, while factorial 01 and 11 were newly trained. These subsequent analyses were not part of the original primary inferential program.

##### S1.6 Historical PSNR–PI Source-Set Provenance

All 44 models retained in Supplementary Table S7 predated the 2026-02-17 peak-loss correction. Thirty-four had nominally nonzero peak-loss weights, but the defective peak-loss component contributed no training gradient; their outputs remain valid evaluations of the models that were actually trained, while the nominal peak-loss axis was inoperative. Separately, nine weight-sweep rows contained wrong evaluation data because a relative split path silently fell back to an unintended three-case split: all eight `plane_mean_max` rows and `global_max_x2p25`. The other seven `global_max` rows were launched in an earlier wave from a working directory in which the fixed split resolved correctly and were unaffected. Table S7 now substitutes the existing corrected six-case reevaluations for those nine rows.

The frozen-package manifest also mapped the six `bio_*` display rows to `flowpi` run identifiers, whereas the printed values came from the corresponding `paper_followup_no_aug_global_max_*` and `paper_followup_no_aug_plane_mean_max_*` runs without `flowpi` in the identifier (seeds 7, 13, and 42). Those six mappings are corrected here. Using the corrected 44 evaluation rows, the historical exploratory Spearman result is $\rho = +0.155$, two-sided $p = 0.315$, $n = 44$. Alternative source reconstructions considered during the audit yielded the same qualitative interpretation, but none is used as evidentiary support in the manuscript.

**Historical execution-window audit.** Retained training logs span 2026-03-07 through 2026-03-12 across the four primary arms. Repository changes touching `src/` or `scripts/` within that interval were launcher, reporting, compatibility, or resume/skip changes that were either unreachable for these fresh runs or behaviorally inert for training and checkpoint selection. The odd-dimension RCAN target-size fallback is reachable for volumes with an odd HR dimension, but its nearest-neighbor fallback change preceded the first retained training timestamp. The historical run directories do not record an executed commit hash, so exact implementation-version identity cannot be independently verified beyond this execution-window audit.

#### Supplementary Table S1: Primary 14-Pair Experiment Matrix

Table S1 lists the 14 seeds used in the primary analysis. All four arms were run for every seed; each cell confirms that one matched run was completed for that seed–arm combination. Detailed experiment records for the primary analysis are preserved in the archived study records and are available from the corresponding author upon reasonable request.

All experiments used: 3D RCAN architecture, 12.65M parameters; 2× downsampling; noise-enabled degradation (SNR [15,30] dB); per-plane mean-of-maxima peak aggregation; no geometric augmentation; 250-epoch cosine annealing learning-rate schedule (peak $10^{-4}$); batch size 1; fixed 12/4/6 stratified train/validation/test split (shared across all runs); checkpoint criterion: validation peak-velocity error; DIPC HPC NVIDIA A100-PCIe (80 GB). In all biomarker arms: $\lambda_{\text{peak}} = 0$ (explicit configuration), contributing no peak-velocity gradient during training.

Supplementary Table S1: Primary 14-pair experiment matrix ($n = 14$ seeds). Each ✓ confirms one completed matched run for that seed–arm combination.

| Seed | Full Biomarker-Aware | Reconstruction-Only | Combined Regularization | Biomarker-Only Control |
|---|---|---|---|---|
| 021 | ✓ | ✓ | ✓ | ✓ |
| 031 | ✓ | ✓ | ✓ | ✓ |
| 037 | ✓ | ✓ | ✓ | ✓ |
| 041 | ✓ | ✓ | ✓ | ✓ |
| 047 | ✓ | ✓ | ✓ | ✓ |
| 053 | ✓ | ✓ | ✓ | ✓ |
| 059 | ✓ | ✓ | ✓ | ✓ |
| 061 | ✓ | ✓ | ✓ | ✓ |
| 067 | ✓ | ✓ | ✓ | ✓ |
| 071 | ✓ | ✓ | ✓ | ✓ |
| 079 | ✓ | ✓ | ✓ | ✓ |
| 083 | ✓ | ✓ | ✓ | ✓ |
| 089 | ✓ | ✓ | ✓ | ✓ |
| 099 | ✓ | ✓ | ✓ | ✓ |

Note: All $\lambda_{\text{peak}} = 0$ in full biomarker-aware and biomarker-only control arms. Combined regularization and biomarker-only control arms used the same 250-epoch schedule, fixed split, and RCAN architecture as the primary pairs.

#### Supplementary Table S2: Per-Seed Four-Arm Results (Primary 14-Pair Analysis)

Table S2 reports per-seed results for all four arms on the primary metrics. All values are MAPE (%) averaged over the fixed 6-case test set. Δ PI (biomarker-aware − reconstruction-only) and Δ PI (biomarker-aware − combined-regularization control) are signed differences; negative values favor the full biomarker-aware arm.

† Seed 047 full biomarker-aware vs. combined-regularization control: both arms selected foundation-phase epoch 35, where all biomarker weights were zero and the active objectives were identical. Their archived validation value, test PI, and DivRMS were exactly equal; the structural tie ($\Delta$PI = 0.0000 pp) was retained.

**Full biomarker-aware vs. reconstruction-only (contextual secondary):** Two-sided Wilcoxon $W_+ = 10$, $p = 0.0052$; one-sided $p = 0.0026$ (sensitivity). Cohen's $d_z = 1.046$ (magnitude), rank-biserial $r = 0.810$. Sign test: 10/14 wins, one-sided $p = 0.090$.

**Full biomarker-aware vs. combined-regularization control (primary benchmark comparison):** Two-sided Wilcoxon $p = 0.0015$; one-sided $p = 0.0007$ (sensitivity). Cohen's $d_z = 1.76$ (magnitude), rank-biserial $r = 1.00$. 13 wins / 1 tie / 0 losses.

**Biomarker-only control vs. reconstruction-only:** Two-sided Wilcoxon $p = 0.0203$. Cohen's $d_z = 0.744$ (magnitude). 10/14 wins.

**Combined regularization vs. reconstruction-only:** Two-sided Wilcoxon $p = 0.0001$. Combined regularization worse in 14/14 seeds.

**Win asymmetry (full biomarker-aware vs. reconstruction-only):** Losses (+0.05, +0.26, +0.51, +0.71 pp) are consistently smaller in magnitude than wins (−0.72 to −3.59 pp).

*Note:* Mean rows are arithmetic means of the 14 per-seed values above. Detailed experiment records for each arm are preserved in the archived study records.

Supplementary Table S2: Per-seed four-arm results ($n = 14$ seeds). PI and DivRMS are MAPE (%). $\Delta$(FBA−RO) = full biomarker-aware minus reconstruction-only; $\Delta$(FBA−combined) = full biomarker-aware minus combined-regularization control; both in pp. Bold $\Delta$ = full biomarker-aware win. Seed 047 combined-regularization control tie marked with †.

| Seed | Full Biomarker-Aware | | Reconstruction-Only | | Combined Regularization | | Biomarker-Only Control | | Δ PI | Δ PI |
|---|---|---|---|---|---|---|---|---|---|---|
| | PI | Div | PI | Div | PI | Div | PI | Div | (FBA−RO) | (FBA−combined) |
| 021 | 13.17 | 4.67 | 15.11 | 7.22 | 17.09 | 5.18 | 15.87 | 7.15 | **−1.94** | **−3.92** |
| 031 | 11.45 | 4.70 | 14.12 | 7.01 | 16.39 | 4.92 | 12.85 | 7.34 | **−2.67** | **−4.94** |
| 037 | 10.90 | 4.78 | 12.95 | 7.01 | 18.40 | 6.56 | 10.52 | 7.48 | **−2.05** | **−7.50** |
| 041 | 11.16 | 4.68 | 14.75 | 7.10 | 19.20 | 6.74 | 14.80 | 7.13 | **−3.59** | **−8.04** |
| 047† | 18.33 | 6.30 | 18.06 | 7.34 | 18.33 | 6.30 | 18.07 | 7.34 | +0.26 | 0.00† |
| 053 | 13.19 | 4.76 | 13.15 | 7.11 | 15.13 | 4.78 | 12.94 | 7.25 | +0.05 | **−1.94** |
| 059 | 11.57 | 4.65 | 14.56 | 7.09 | 14.96 | 4.93 | 11.23 | 7.42 | **−2.99** | **−3.39** |
| 061 | 10.77 | 4.69 | 13.02 | 7.04 | 16.49 | 4.73 | 9.77 | 7.36 | **−2.25** | **−5.72** |
| 067 | 13.45 | 4.64 | 12.93 | 7.18 | 17.95 | 4.94 | 14.38 | 7.39 | +0.51 | **−4.50** |
| 071 | 17.33 | 6.08 | 16.62 | 7.35 | 17.35 | 6.08 | 16.52 | 7.38 | +0.71 | **−0.02** |
| 079 | 12.46 | 4.75 | 14.32 | 6.85 | 16.00 | 4.76 | 13.40 | 7.02 | **−1.87** | **−3.54** |
| 083 | 11.85 | 4.80 | 13.20 | 6.96 | 17.88 | 4.96 | 11.09 | 7.37 | **−1.35** | **−6.03** |
| 089 | 12.34 | 4.90 | 13.07 | 7.08 | 17.04 | 4.78 | 11.10 | 7.46 | **−0.72** | **−4.70** |
| 099 | 10.72 | 4.77 | 13.16 | 7.07 | 17.40 | 4.75 | 11.07 | 7.51 | **−2.44** | **−6.68** |
| **Mean** | 12.76 | 4.94 | 14.22 | 7.10 | 17.12 | 5.31 | 13.12 | 7.33 | **−1.45** | **−4.35** |
| **SD** | 2.34 | 0.54 | 1.55 | 0.14 | 1.23 | 0.75 | 2.51 | 0.14 | 1.39 | 2.47 |

#### Supplementary Table S3: Validation-Criterion Rerun Comparison — All Four RCAN Arms, Full 14-Seed Per-Seed Detail

Table S3 reports per-seed PI values from the primary peak-velocity-validation runs and from matched reruns using PI as the validation / early-stopping criterion for all 14 seeds and all four RCAN arms. PI-validation reruns were completed for the full biomarker-aware and reconstruction-only arms (five initial seeds 031, 037, 041, 047, 083 combined with nine additional seeds 021, 053, 059, 061, 067, 071, 079, 089, 099) and for the combined-regularization control and biomarker-only control arms (full 14-seed runs completed as a subsequent extension). $\Delta$ = PI from the PI-validation rerun − PI from the corresponding primary run; negative values indicate the PI-validation rerun is better.

**Summary statistics (all four arms):**

- **Full biomarker-aware:** mean $\Delta = -1.76$ pp (95% CI [−3.00, −0.76]); two-sided $p = 0.0004$; W/T/L = 13/0/1.
- **Reconstruction-only:** mean $\Delta = -1.24$ pp (95% CI [−2.07, −0.54]); two-sided $p = 0.0001$; W/T/L = 14/0/0.
- **Combined regularization:** mean $\Delta = -2.23$ pp (95% CI [−3.01, −1.48]); two-sided $p < 0.0001$; W/T/L = 14/0/0.
- **Biomarker-only control:** mean $\Delta = -2.46$ pp (95% CI [−3.67, −1.34]); two-sided $p = 0.0006$; W/T/L = 13/0/1.

All $p$-values from two-sided paired Wilcoxon signed-rank test; 95% CIs from bootstrap resampling (1000 resamples, percentile interval). The single biomarker-only control loss (seed 061, +0.58 pp) occurs in the seed with the lowest peak-velocity-aligned PI in that arm (9.77%, already well below the 15% engineering reference); this seed's PI-aligned checkpoint is marginally worse but the absolute PI value remains low. All four arms show consistent directional improvement, supporting the interpretation that using PI rather than peak velocity as the validation criterion materially affects endpoint estimates in this benchmark regardless of training loss.

*Note:* Peak-velocity-aligned values are from the primary experiment families (Supplementary Table S2). PI-aligned values are from dedicated matched reruns of each arm using the same training configuration, with only the validation / early-stopping criterion changed from peak velocity to PI validation error. For the full biomarker-aware and reconstruction-only arms, five seeds were completed as an initial subset (seeds 031, 037, 041, 047, 083); the remaining nine seeds were added subsequently. For the combined-regularization control and biomarker-only control arms, all 14 PI-aligned runs were conducted as a subsequent extension. Detailed records for all PI-aligned runs are preserved in the archived study records. Bootstrap CIs used 1000 resamples (percentile interval); note that the primary benchmark comparison used 10,000 resamples (main text, Section 2.6).

Supplementary Table S3: Per-seed matched-rerun comparison for all four RCAN arms ($n = 14$ seeds each). "Peak-velocity-aligned" = checkpoint selected in the primary run by minimum validation peak-velocity error; "PI-aligned" = checkpoint selected in the matched rerun by minimum validation PI error. $\Delta$ = PI-aligned − peak-velocity-aligned (pp); negative values favor the PI-validation rerun. Bold $\Delta$ = PI-validation rerun win. All peak-velocity-aligned values are from primary experiment families; PI-aligned values are from dedicated matched PI-validation reruns. Seed 061 biomarker-only control loss (+0.58 pp) is noted; that seed's peak-velocity-aligned PI (9.77%) was already below the arm mean.

**Panel A. Full biomarker-aware and reconstruction-only arms.**

| | Full Biomarker-Aware | | | Reconstruction-Only | | |
|---|---|---|---|---|---|---|
| Seed | Peak-velocity aligned PI(%) | PI aligned PI(%) | $\Delta$ (pp) | Peak-velocity aligned PI(%) | PI aligned PI(%) | $\Delta$ (pp) |
| 021 | 13.17 | 10.63 | **−2.54** | 15.11 | 13.24 | **−1.87** |
| 031 | 11.45 | 11.31 | **−0.14** | 14.12 | 13.33 | **−0.78** |
| 037 | 10.90 | 10.66 | **−0.24** | 12.95 | 12.70 | **−0.25** |
| 041 | 11.16 | 10.93 | **−0.23** | 14.75 | 13.28 | **−1.47** |
| 047 | 18.33 | 11.44 | **−6.89** | 18.06 | 13.23 | **−4.83** |
| 053 | 13.19 | 11.24 | **−1.95** | 13.15 | 12.64 | **−0.50** |
| 059 | 11.57 | 10.85 | **−0.72** | 14.56 | 13.25 | **−1.31** |
| 061 | 10.77 | 10.72 | **−0.06** | 13.02 | 12.79 | **−0.23** |
| 067 | 13.45 | 10.80 | **−2.64** | 12.93 | 12.86 | **−0.07** |
| 071 | 17.33 | 10.87 | **−6.46** | 16.62 | 12.55 | **−4.08** |
| 079 | 12.46 | 11.25 | **−1.21** | 14.32 | 13.13 | **−1.20** |
| 083 | 11.85 | 11.54 | **−0.31** | 13.20 | 13.14 | **−0.06** |
| 089 | 12.34 | 11.07 | **−1.27** | 13.07 | 12.78 | **−0.28** |
| 099 | 10.72 | 10.81 | +0.09 | 13.16 | 12.72 | **−0.44** |
| Mean | 12.76 | 11.01 | −1.76 | 14.22 | 12.97 | −1.24 |
| SD | 2.34 | 0.29 | 2.19 | 1.55 | 0.27 | 1.43 |

**Panel B. Combined regularization and biomarker-only control arms.**

| | Combined Regularization | | | Biomarker-Only Control | | |
|---|---|---|---|---|---|---|
| Seed | Peak-velocity aligned PI(%) | PI aligned PI(%) | $\Delta$ (pp) | Peak-velocity aligned PI(%) | PI aligned PI(%) | $\Delta$ (pp) |
| 021 | 17.09 | 14.96 | **−2.12** | 15.87 | 11.38 | **−4.49** |
| 031 | 16.39 | 16.09 | **−0.30** | 12.85 | 10.87 | **−1.98** |
| 037 | 18.40 | 14.31 | **−4.09** | 10.52 | 10.20 | **−0.33** |
| 041 | 19.20 | 14.15 | **−5.05** | 14.80 | 10.77 | **−4.02** |
| 047 | 18.33 | 15.08 | **−3.24** | 18.07 | 10.93 | **−7.14** |
| 053 | 15.13 | 14.77 | **−0.36** | 12.94 | 10.93 | **−2.01** |
| 059 | 14.96 | 14.23 | **−0.74** | 11.23 | 10.32 | **−0.92** |
| 061 | 16.49 | 15.63 | **−0.86** | 9.77 | 10.35 | +0.58 |
| 067 | 17.95 | 14.58 | **−3.37** | 14.38 | 10.46 | **−3.93** |
| 071 | 17.35 | 15.14 | **−2.21** | 16.52 | 10.80 | **−5.72** |
| 079 | 16.00 | 15.42 | **−0.58** | 13.40 | 10.71 | **−2.69** |
| 083 | 17.88 | 14.58 | **−3.30** | 11.09 | 10.69 | **−0.40** |
| 089 | 17.04 | 14.54 | **−2.50** | 11.10 | 10.37 | **−0.72** |
| 099 | 17.40 | 14.88 | **−2.52** | 11.07 | 10.42 | **−0.65** |
| Mean | 17.12 | 14.88 | −2.23 | 13.12 | 10.66 | −2.46 |
| SD | 1.23 | 0.56 | 1.50 | 2.51 | 0.32 | 2.29 |

#### Supplementary Table S4: TemporalUNet4D Probe — Full 5-Seed Results

Table S4 reports per-seed TUNet results for all five seeds and four arms. Seed 047 was unstable in the full biomarker-aware and biomarker-only control arms and is excluded symmetrically from all four arms in the four-seed summary. Seed 031 biomarker-only control showed elevated DivRMS and lower PSNR. The four-seed summary (seeds 031, 037, 041, 083; seed 047 excluded symmetrically from all four arms) is shown in the main text; the full 5-seed table is provided here for transparency.

Supplementary Table S4: TemporalUNet4D full 5-seed four-arm results. PI and DivRMS are MAPE (%). PSNR in dB. Flags: ‡ = unstable run (seed 047 full biomarker-aware and biomarker-only control arms, excluded symmetrically from all four arms in the four-seed summary); $*$ = elevated DivRMS / lower PSNR (seed 031 biomarker-only control, retained in four-seed summary with footnote).

| Seed | Arm | PI (%) | DivRMS (%) | PSNR (dB) |
|---|---|---|---|---|
| 031 | Reconstruction-only | 10.51 | 9.86 | 37.28 |
| | Combined regularization | 17.93 | 6.64 | 37.88 |
| | Biomarker-only control* | 13.70 | 17.94 | 32.46 |
| | Full biomarker-aware | 17.93 | 7.43 | 37.68 |
| 037 | Reconstruction-only | 10.26 | 8.87 | 38.12 |
| | Combined regularization | 18.04 | 6.65 | 37.70 |
| | Biomarker-only control | 12.68 | 7.98 | 36.44 |
| | Full biomarker-aware | 18.22 | 7.21 | 38.00 |
| 041 | Reconstruction-only | 14.18 | 9.33 | 37.29 |
| | Combined regularization | 19.61 | 6.63 | 37.56 |
| | Biomarker-only control | 12.97 | 8.37 | 36.16 |
| | Full biomarker-aware | 18.82 | 6.96 | 37.70 |
| 047 | Reconstruction-only | 14.48 | 9.73 | 36.79 |
| | Combined regularization | 17.88 | 6.72 | 37.54 |
| | Biomarker-only control‡ | unstable | – | – |
| | Full biomarker-aware‡ | unstable | – | – |
| 083 | Reconstruction-only | 14.25 | 8.98 | 36.79 |
| | Combined regularization | 18.66 | 6.48 | 37.74 |
| | Biomarker-only control | 13.25 | 8.90 | 36.90 |
| | Full biomarker-aware | 17.75 | 6.28 | 37.27 |
| **Four-seed summary (seeds 031, 037, 041, 083; seed 047 excluded symmetrically from all arms):** | | | | |
| Mean ± SD | Reconstruction-only | 12.30 ± 2.21 | 9.26 ± 0.45 | 37.37 ± 0.55 |
| | Combined regularization | 18.56 ± 0.77 | 6.60 ± 0.08 | 37.72 ± 0.13 |
| | Biomarker-only control* | 13.15 ± 0.43 | 10.80 ± 4.78 | 35.49 ± 2.04 |
| | Full biomarker-aware | 18.18 ± 0.47 | 6.97 ± 0.50 | 37.66 ± 0.30 |

*Seed 031 biomarker-only control DivRMS = 17.94%, PSNR = 32.46 dB are substantially elevated/degraded relative to the other three seeds; this run is retained in the four-seed summary but inflates the biomarker-only control SD. The elevated DivRMS in this run may reflect a training instability in the absence of combined regularization for this seed.

‡Seed 047 full biomarker-aware and biomarker-only control arms were classified as unstable (training divergence or extreme metric values) and are excluded symmetrically from all four arms in the four-seed summary. The two-arm data (reconstruction-only, combined-regularization control) for seed 047 are retained here for completeness.

**Directional interpretation:** Within TUNet, the arms containing the historical combined divergence–temporal formulation consistently improved DivRMS relative to reconstruction-only in the four-seed summary, but also showed higher PI. This probe does not isolate divergence from temporal smoothing and should not be interpreted as a head-to-head RCAN-vs-TUNet benchmark.

#### Supplementary Table S5: Harder-Noise Robustness Screen — Full 5-Seed Results

Table S5 reports per-seed harder-noise results for all five seeds and four RCAN arms (SNR [10,20] dB; seeds 021, 031, 041, 047, 071). The five-seed summary block reproduces the main-manuscript harder-noise summary and adds PSNR SD because full per-seed values are available here. This is directional/descriptive evidence only and does not constitute a second primary study.

Supplementary Table S5: Harder-noise robustness screen: full 5-seed four-arm results. PI and DivRMS are MAPE (%); PSNR in dB. The five-seed summary block gives mean ± SD for all three metrics. This table is descriptive only; no inferential statistics are reported.

| Seed | Arm | PI (%) | DivRMS (%) | PSNR (dB) |
|---|---|---|---|---|
| 021 | Reconstruction-only | 11.63 | 7.85 | 39.36 |
| | Combined regularization | 18.26 | 6.00 | 38.65 |
| | Biomarker-only control | 9.97 | 8.15 | 38.95 |
| | Full biomarker-aware | 17.24 | 5.82 | 38.52 |
| 031 | Reconstruction-only | 11.83 | 7.89 | 39.25 |
| | Combined regularization | 18.97 | 6.46 | 38.44 |
| | Biomarker-only control | 11.07 | 8.10 | 38.93 |
| | Full biomarker-aware | 11.51 | 6.08 | 38.46 |
| 041 | Reconstruction-only | 12.78 | 7.77 | 39.34 |
| | Combined regularization | 16.84 | 5.90 | 39.07 |
| | Biomarker-only control | 9.44 | 8.13 | 39.02 |
| | Full biomarker-aware | 11.53 | 6.03 | 38.73 |
| 047 | Reconstruction-only | 13.85 | 8.03 | 38.87 |
| | Combined regularization | 17.43 | 6.06 | 38.67 |
| | Biomarker-only control | 8.99 | 8.24 | 38.86 |
| | Full biomarker-aware | 17.42 | 6.26 | 38.66 |
| 071 | Reconstruction-only | 17.08 | 8.30 | 38.33 |
| | Combined regularization | 18.03 | 7.06 | 38.34 |
| | Biomarker-only control | 11.44 | 7.93 | 39.02 |
| | Full biomarker-aware | 14.21 | 5.91 | 38.86 |
| **Five-seed summary (descriptive only):** | | | | |
| Mean ± SD | Reconstruction-only | 13.43 ± 2.22 | 7.97 ± 0.21 | 39.03 ± 0.44 |
| | Combined regularization | 17.91 ± 0.81 | 6.30 ± 0.48 | 38.63 ± 0.28 |
| | Biomarker-only control | 10.18 ± 1.05 | 8.11 ± 0.11 | 38.96 ± 0.07 |
| | Full biomarker-aware | 14.38 ± 2.91 | 6.02 ± 0.17 | 38.65 ± 0.16 |

**Directional interpretation:** Within this five-seed screen, the historical combined-control arm had higher PI and lower DivRMS than reconstruction-only in 5/5 seeds; this does not identify which regularization component produced the PI change. Biomarker-only control improved PI relative to reconstruction-only in 5/5 seeds. Full biomarker-aware vs. reconstruction-only was mixed (3/5 wins; mean +0.95 pp favoring reconstruction-only). These are directional scope-bounding results only.

#### Supplementary Table S6: Noise-Enabled vs. Noise-Free Degradation Controls

Table S6 reports the eight matched standard-degradation control runs referred to in main-text Section 3.5. All runs use RCAN, seed 042, matched training settings, and peak-velocity validation; field-by-field comparison of the archived configurations confirmed that training, loss, scheduling, and checkpoint settings were identical within each pair, with only the intended additive-noise toggle and descriptive run identifiers differing. These runs used historical biomarker-optimized configurations with nominally nonzero peak-velocity, percentile-velocity, flow-rate, and pulsatility weights. Because they predated the documented peak-gradient correction, the defective peak-loss component contributed no training gradient and the nominal peak-loss axis was inoperative; the other biomarker-loss components were active. Both members of each pair were trained under the same contemporaneous peak-loss implementation, so the comparison remains internally matched with respect to the training configuration and that implementation. It does not establish what the numerical contrast would have been under a functioning peak-velocity loss. This table provides supporting contextual evidence only and should not be interpreted as mechanistic proof.

Supplementary Table S6: Eight matched standard-degradation control runs from a single matched seed (seed 042) comparing noise-enabled and noise-free degradation. Metrics are MAPE (%) except PSNR (dB). Values are reproduced directly from the archived runs. A separate `psf0.0` noise-free sharp-control variant, distinct from the `psf0.0` standard-degradation row listed here, is excluded because it falls outside the matched standard-degradation on/off comparison discussed in the manuscript.

| PSF setting | Noise | Peak velocity (%) | Flow rate (%) | PI (%) | PSNR (dB) |
|---|---|---|---|---|---|
| `psf0.4` | enabled | 5.71 | 5.13 | 15.66 | 40.10 |
| `psf0.4` | noise-free | 26.31 | 26.66 | 15.91 | 34.56 |
| `psf0.2` | enabled | 4.70 | 4.95 | 10.58 | 40.28 |
| `psf0.2` | noise-free | 23.72 | 24.25 | 15.78 | 35.18 |
| `psf0.1` | enabled | 4.61 | 4.91 | 10.62 | 40.27 |
| `psf0.1` | noise-free | 23.27 | 23.89 | 15.75 | 35.27 |
| `psf0.0` | enabled | 4.65 | 4.85 | 11.00 | 40.19 |
| `psf0.0`$^\dagger$ | noise-free | 23.27 | 23.89 | 15.75 | 35.27 |

$^\dagger$ The archived seed-042 standard-degradation outputs for the `psf0.0` and `psf0.1` noise-free rows differ slightly at higher precision, but all four displayed metrics round to the same two-decimal table values. We therefore retain the displayed values as rounded archival summaries rather than inferring a larger distinction.

Within each matched pair, removing noise increased peak-velocity error by 18.62–20.60 pp and flow-rate error by 18.98–21.53 pp. PI changed little at `psf0.4` (+0.25 pp) but worsened by 4.75–5.21 pp in the `psf0.0`–`psf0.2` pairs; the upper endpoint is calculated from the unrounded archived values. These are single-seed, benchmark-specific contextual observations and should not be interpreted as mechanistic or inferential evidence.

#### Supplementary Table S7: Historical Exploratory PSNR–PI Source Data

Table S7 retains the audited 44-run standard-degradation RCAN screening set as historical exploratory material. Nine rows now use their existing corrected six-case reevaluations and six display-to-run mappings have been corrected, as detailed in Supplementary Note S1.6. This source set is not evidentiary support for the conclusions. Supplementary Figure S1 plots the corrected rows.

Supplementary Table S7: Historical exploratory PSNR–PI source data with corrected evaluation provenance.

| Run identifier | PkVel (%) | FlRate (%) | PI (%) | PSNR (dB) |
|---|---|---|---|---|
| **Matched comparison runs (22 runs)** | | | | |
| `matched_allup_gm_x1p75_s13` | 6.02 | 4.56 | 11.60 | 39.60 |
| `matched_allup_gm_x1p75_s21` | 4.65 | 4.72 | 10.96 | 39.93 |
| `matched_allup_gm_x1p75_s7` | 5.61 | 5.66 | 12.25 | 39.44 |
| `matched_allup_gm_x1p75_s99` | 4.76 | 3.58 | 10.69 | 39.77 |
| `matched_allup_pm_x1p5_s13` | 5.03 | 5.33 | 10.52 | 39.87 |
| `matched_allup_pm_x1p5_s21` | 5.38 | 4.67 | 10.62 | 39.90 |
| `matched_allup_pm_x1p5_s7` | 9.76 | 6.07 | 17.87 | 39.34 |
| `matched_allup_pm_x1p5_s99` | 6.52 | 3.12 | 11.26 | 39.64 |
| `matched_bio_no_aug_gm_s21` | 4.96 | 5.46 | 14.09 | 40.09 |
| `matched_bio_no_aug_gm_s99` | 5.41 | 4.25 | 11.78 | 40.09 |
| `matched_bio_no_aug_pm_s21` | 4.59 | 5.34 | 14.05 | 40.18 |
| `matched_bio_no_aug_pm_s99` | 5.01 | 3.93 | 12.11 | 39.98 |
| `matched_recon_no_aug_gm_s13` | 5.10 | 3.98 | 14.79 | 40.06 |
| `matched_recon_no_aug_gm_s21` | 4.05 | 4.12 | 13.19 | 40.51 |
| `matched_recon_no_aug_gm_s42` | 5.18 | 5.15 | 15.12 | 40.29 |
| `matched_recon_no_aug_gm_s7` | 5.91 | 3.53 | 14.32 | 39.94 |
| `matched_recon_no_aug_gm_s99` | 4.77 | 3.92 | 13.46 | 40.48 |
| `matched_recon_no_aug_pm_s13` | 5.09 | 3.97 | 14.79 | 40.06 |
| `matched_recon_no_aug_pm_s21` | 4.04 | 4.12 | 13.19 | 40.51 |
| `matched_recon_no_aug_pm_s42` | 5.19 | 5.14 | 15.14 | 40.29 |
| `matched_recon_no_aug_pm_s7` | 5.90 | 3.53 | 14.32 | 39.94 |
| `matched_recon_no_aug_pm_s99` | 4.77 | 3.91 | 13.46 | 40.48 |
| **Biomarker follow-up runs (6 runs)** | | | | |
| `bio_global_max_s13` | 5.29 | 5.05 | 16.47 | 39.84 |
| `bio_global_max_s42` | 7.46 | 7.96 | 18.80 | 39.49 |
| `bio_global_max_s7` | 4.70 | 4.95 | 12.55 | 39.86 |
| `bio_plane_mean_max_s13` | 4.69 | 4.97 | 12.36 | 40.01 |
| `bio_plane_mean_max_s42` | 4.93 | 5.01 | 12.01 | 40.04 |
| `bio_plane_mean_max_s7` | 4.83 | 4.58 | 12.76 | 39.78 |
| **Weight-sweep runs (16 runs)** | | | | |
| `allup_global_max_x1p0` | 6.15 | 5.15 | 15.94 | 40.07 |
| `allup_global_max_x1p25` | 4.61 | 4.80 | 12.13 | 39.77 |
| `allup_global_max_x1p4` | 4.93 | 4.18 | 10.95 | 39.89 |
| `allup_global_max_x1p5` | 4.87 | 4.89 | 14.30 | 39.99 |
| `allup_global_max_x1p6` | 5.18 | 5.55 | 15.29 | 39.99 |
| `allup_global_max_x1p75` | 4.35 | 4.42 | 10.82 | 39.76 |
| `allup_global_max_x2p0` | 4.16 | 6.13 | 13.11 | 39.65 |
| `allup_global_max_x2p25` | 4.41 | 4.20 | 13.50 | 39.87 |
| `allup_plane_mean_max_x1p0` | 7.71 | 8.81 | 19.77 | 39.41 |
| `allup_plane_mean_max_x1p25` | 4.55 | 5.42 | 10.73 | 39.86 |
| `allup_plane_mean_max_x1p4` | 5.74 | 4.92 | 15.24 | 40.09 |
| `allup_plane_mean_max_x1p5` | 4.47 | 4.61 | 11.10 | 39.82 |
| `allup_plane_mean_max_x1p6` | 4.74 | 5.12 | 10.67 | 39.87 |
| `allup_plane_mean_max_x1p75` | 6.20 | 5.48 | 16.68 | 40.03 |
| `allup_plane_mean_max_x2p0` | 7.18 | 8.36 | 18.93 | 39.51 |
| `allup_plane_mean_max_x2p25` | 6.15 | 5.14 | 16.16 | 40.12 |

*Note:* These 44 rows are retained as a corrected historical audit trail and are plotted in Supplementary Figure S1; they are not used to support the conclusions. Run identifiers preserve the original display labels. The prefixes `matched`, `bio`, and `allup` denote the three configuration families. The labels `gm` and `global_max` denote global-maximum peak estimation, whereas `pm` and `plane_mean_max` denote plane-wise mean-maximum peak estimation. In the weight-sweep identifiers, the number following `x` gives the relative weight setting and the number following `s` gives the seed; `no_aug` denotes no geometric augmentation. Corrected run mappings and the distinction between wrong evaluation data and an inoperative nominal peak-loss axis are given in Supplementary Note S1.6.

#### Supplementary Table S8: Experiment-Family Provenance Summary

Supplementary Table S8: Experiment-family provenance and evidentiary roles. CR denotes the combined divergence–temporal-regularization control. Archived identifiers may retain the historical `physctl` string; that identifier should not be interpreted as divergence regularization alone. The component and factorial analyses were performed after the original primary analysis. The table additionally includes the unnumbered historical PSNR–PI source audit for provenance; it is not one of the eight declared experiment families.

| Family | Seed / sample base | Arms | Model | Validation | Evidence tier | Detailed reporting |
|---|---|---|---|---|---|---|
| Primary 4-arm paired analysis | 14 matched seeds | RO, CR, BIO, FBA | RCAN | Peak velocity | Primary | Main Results §3.1; Supplementary Tables S1–S2 and S9 |
| Three-level ablation | 3 seeds | L1–L3 | RCAN | Peak velocity | Directional | Supplementary Figure S2 |
| PI-validation matched-rerun study | Same 14 seeds | RO, CR, BIO, FBA | RCAN | PI | Contextual | Main Results §3.4; Supplementary Table S3 |
| Noise-free degradation comparison | 8 matched runs (seed 042) | Historical biomarker-optimized configurations | RCAN | Peak velocity | Descriptive | Main Results §3.5; Supplementary Table S6 |
| Harder-noise robustness screen | 5 seeds | RO, CR, BIO, FBA | RCAN | Peak velocity | Directional | Main Results §3.6; Supplementary Table S5 |
| TemporalUNet4D probe | 5 seeds (4-seed summary) | RO, CR, BIO, FBA | TUNet | Peak velocity | Directional | Main Results §3.7; Supplementary Table S4 |
| Historical exploratory PSNR–PI source audit | 44 pre-correction models in Supplementary Table S7 and Supplementary Figure S1 | — | RCAN | — | Non-evidentiary audit trail | Supplementary Note S1.6, Table S7, and Figure S1 |
| Regularization-component ablation | Same 14 seeds | R, RD, RT, RDT | RCAN | Peak velocity | Additional controlled; shares R/RDT with factorial | Main Results §3.2; Supplementary Tables S10–S11 |
| Matched 2×2 factorial | Same 14 seeds | 00, 10, 01, 11 | RCAN | Peak velocity | Additional controlled; 00=R and 10=RDT | Main Results §3.3; Supplementary Tables S10–S13 |

#### Supplementary Table S9: Per-Case PI Consistency (Primary Benchmark Comparison)

Table S9 reports per-case PI results for the primary benchmark comparison (full biomarker-aware vs. combined-regularization control) across all 6 test cases and 14 seeds. For each case, mean ± SD across 14 seeds and the paired difference Δ (full biomarker-aware − combined-regularization control, mean) are shown. Win count = number of seeds where full biomarker-aware PI < combined-regularization control PI. This table characterizes whether the primary PI improvement is distributed across all test cases or concentrated in a subset; it does not characterize patient-level generalization.

Supplementary Table S9: Per-case PI consistency for the primary benchmark comparison (full biomarker-aware vs. combined-regularization control; $n = 14$ seeds per case). PI values are MAPE (%); Δ = full biomarker-aware − combined-regularization control mean (pp); negative values favor the full biomarker-aware arm. Win = number of seeds with full biomarker-aware PI < combined-regularization control PI.

| Case | FBA mean | FBA SD | Combined-reg. mean | Combined-reg. SD | Δ mean | Win |
|---|---|---|---|---|---|---|
| C1 | 20.92 | 2.42 | 24.86 | 0.76 | −3.95 | 13/14 |
| C2 | 13.27 | 2.51 | 17.43 | 1.00 | −4.16 | 12/14 |
| C3 | 9.18 | 4.28 | 17.06 | 3.37 | −7.87 | 13/14 |
| C4 | 10.13 | 2.62 | 14.55 | 0.83 | −4.43 | 13/14 |
| C5 | 19.13 | 1.67 | 22.13 | 0.59 | −3.00 | 12/14 |
| C6 | 3.96 | 1.67 | 6.66 | 1.51 | −2.70 | 13/14 |
| **Mean** | **12.76** | – | **17.12** | – | **−4.35** | – |

The full biomarker-aware arm showed lower PI than the combined-regularization control arm in at least 12 of 14 seeds for every test case (range: 12–13/14 wins per case), indicating that the primary PI improvement is distributed across all six test cases rather than being concentrated in a single outlier case. The mean Δ ranges from −2.70 pp (case C6, lowest absolute PI) to −7.87 pp (case C3); the wider range and SD in C3 reflect greater seed-to-seed variability in that case. The per-case mean rows aggregate to the reported arm-level means: full biomarker-aware 12.76% and combined-regularization control 17.12%.

#### Supplementary Table S10: Full Arm Summaries for Additional Controlled Analyses

Values summarize the 14 matched training seeds evaluated on the same fixed six-case test set.

| Analysis | Arm | Metric | Mean | SD | Median |
|---|---|---|---|---|---|
| Component | R | PI error (%) | 15.045 | 1.440 | 14.620 |
| Component | R | Flow-rate error (%) | 5.384 | 0.761 | 5.230 |
| Component | R | Peak-velocity error (%) | 6.351 | 1.267 | 5.842 |
| Component | R | DivRMS (%) | 7.123 | 0.142 | 7.098 |
| Component | R | PSNR (dB) | 39.810 | 0.304 | 39.911 |
| Component | R | SSIM | 0.826 | 0.007 | 0.828 |
| Component | R | NMSE | 0.050 | 0.004 | 0.048 |
| Component | RD | PI error (%) | 15.079 | 1.426 | 14.444 |
| Component | RD | Flow-rate error (%) | 5.182 | 0.865 | 5.019 |
| Component | RD | Peak-velocity error (%) | 6.063 | 1.307 | 5.569 |
| Component | RD | DivRMS (%) | 5.254 | 0.442 | 5.140 |
| Component | RD | PSNR (dB) | 39.776 | 0.260 | 39.856 |
| Component | RD | SSIM | 0.820 | 0.006 | 0.821 |
| Component | RD | NMSE | 0.050 | 0.003 | 0.049 |
| Component | RDT | PI error (%) | 17.259 | 1.258 | 17.344 |
| Component | RDT | Flow-rate error (%) | 6.193 | 1.042 | 6.203 |
| Component | RDT | Peak-velocity error (%) | 6.963 | 1.974 | 6.384 |
| Component | RDT | DivRMS (%) | 5.349 | 0.740 | 4.951 |
| Component | RDT | PSNR (dB) | 39.579 | 0.388 | 39.690 |
| Component | RDT | SSIM | 0.826 | 0.008 | 0.825 |
| Component | RDT | NMSE | 0.052 | 0.005 | 0.051 |
| Component | RT | PI error (%) | 16.607 | 1.374 | 16.507 |
| Component | RT | Flow-rate error (%) | 6.095 | 1.002 | 5.820 |
| Component | RT | Peak-velocity error (%) | 6.477 | 2.211 | 5.744 |
| Component | RT | DivRMS (%) | 6.693 | 0.395 | 6.559 |
| Component | RT | PSNR (dB) | 39.721 | 0.430 | 39.808 |
| Component | RT | SSIM | 0.833 | 0.011 | 0.837 |
| Component | RT | NMSE | 0.051 | 0.005 | 0.049 |
| Factorial | 00 | PI error (%) | 15.045 | 1.440 | 14.620 |
| Factorial | 00 | Flow-rate error (%) | 5.384 | 0.761 | 5.230 |
| Factorial | 00 | Peak-velocity error (%) | 6.351 | 1.267 | 5.842 |
| Factorial | 00 | DivRMS (%) | 7.123 | 0.142 | 7.098 |
| Factorial | 00 | PSNR (dB) | 39.810 | 0.304 | 39.911 |
| Factorial | 00 | SSIM | 0.826 | 0.007 | 0.828 |
| Factorial | 00 | NMSE | 0.050 | 0.004 | 0.048 |
| Factorial | 01 | PI error (%) | 12.809 | 2.342 | 12.151 |
| Factorial | 01 | Flow-rate error (%) | 5.478 | 0.635 | 5.602 |
| Factorial | 01 | Peak-velocity error (%) | 6.242 | 0.916 | 6.307 |
| Factorial | 01 | DivRMS (%) | 7.295 | 0.195 | 7.318 |
| Factorial | 01 | PSNR (dB) | 39.752 | 0.224 | 39.703 |
| Factorial | 01 | SSIM | 0.817 | 0.010 | 0.817 |
| Factorial | 01 | NMSE | 0.051 | 0.003 | 0.051 |
| Factorial | 10 | PI error (%) | 17.259 | 1.258 | 17.344 |
| Factorial | 10 | Flow-rate error (%) | 6.193 | 1.042 | 6.203 |
| Factorial | 10 | Peak-velocity error (%) | 6.963 | 1.974 | 6.384 |
| Factorial | 10 | DivRMS (%) | 5.349 | 0.740 | 4.951 |
| Factorial | 10 | PSNR (dB) | 39.579 | 0.388 | 39.690 |
| Factorial | 10 | SSIM | 0.826 | 0.008 | 0.825 |
| Factorial | 10 | NMSE | 0.052 | 0.005 | 0.051 |
| Factorial | 11 | PI error (%) | 13.134 | 2.569 | 12.038 |
| Factorial | 11 | Flow-rate error (%) | 5.482 | 0.565 | 5.388 |
| Factorial | 11 | Peak-velocity error (%) | 5.649 | 1.223 | 5.278 |
| Factorial | 11 | DivRMS (%) | 4.947 | 0.530 | 4.755 |
| Factorial | 11 | PSNR (dB) | 39.724 | 0.207 | 39.767 |
| Factorial | 11 | SSIM | 0.819 | 0.009 | 0.819 |
| Factorial | 11 | NMSE | 0.051 | 0.003 | 0.050 |

#### Supplementary Table S11: Full Paired Comparisons

Differences are first arm minus second arm. For error metrics and DivRMS, a negative difference is favorable; for PSNR and SSIM, a positive difference is favorable. W/T/L follows the favorable direction for each metric. Confidence intervals are percentile-bootstrap intervals and $p$ values are from two-sided paired Wilcoxon tests.

| Family | Contrast | Metric | Mean diff. | 95% CI | $p$ | W/T/L |
|---|---|---|---|---|---|---|
| Comp. | RD-vs-R | PI error (%) | 0.034 | [-0.705, 0.809] | 0.6257 | 5/0/9 |
| Comp. | RD-vs-R | Flow-rate error (%) | -0.202 | [-0.656, 0.259] | 0.3575 | 8/0/6 |
| Comp. | RD-vs-R | Peak-velocity error (%) | -0.288 | [-0.610, 0.124] | 0.02026 | 12/0/2 |
| Comp. | RD-vs-R | DivRMS (%) | -1.869 | [-2.025, -1.676] | 0.0001221 | 14/0/0 |
| Comp. | RD-vs-R | PSNR (dB) | -0.033 | [-0.146, 0.068] | 0.7609 | 7/0/7 |
| Comp. | RD-vs-R | SSIM | -0.006 | [-0.009, -0.003] | 0.004028 | 3/0/11 |
| Comp. | RD-vs-R | NMSE | 0.000 | [-0.001, 0.002] | 0.8077 | 6/0/8 |
| Comp. | RT-vs-R | PI error (%) | 1.562 | [0.594, 2.439] | 0.0131 | 1/1/12 |
| Comp. | RT-vs-R | Flow-rate error (%) | 0.710 | [0.170, 1.307] | 0.03305 | 2/1/11 |
| Comp. | RT-vs-R | Peak-velocity error (%) | 0.126 | [-0.665, 1.188] | 0.3824 | 10/1/3 |
| Comp. | RT-vs-R | DivRMS (%) | -0.430 | [-0.583, -0.253] | 0.002977 | 12/1/1 |
| Comp. | RT-vs-R | PSNR (dB) | -0.089 | [-0.298, 0.088] | 0.9165 | 7/1/6 |
| Comp. | RT-vs-R | SSIM | 0.007 | [0.003, 0.010] | 0.007132 | 11/1/2 |
| Comp. | RT-vs-R | NMSE | 0.001 | [-0.001, 0.003] | 0.8613 | 7/1/6 |
| Comp. | RDT-vs-R | PI error (%) | 2.214 | [1.541, 2.916] | 0.0001221 | 0/0/14 |
| Comp. | RDT-vs-R | Flow-rate error (%) | 0.809 | [0.340, 1.351] | 0.005249 | 3/0/11 |
| Comp. | RDT-vs-R | Peak-velocity error (%) | 0.612 | [-0.106, 1.566] | 0.3258 | 7/0/7 |
| Comp. | RDT-vs-R | DivRMS (%) | -1.774 | [-2.096, -1.402] | 0.0001221 | 14/0/0 |
| Comp. | RDT-vs-R | PSNR (dB) | -0.230 | [-0.412, -0.072] | 0.05798 | 4/0/10 |
| Comp. | RDT-vs-R | SSIM | -0.000 | [-0.003, 0.002] | 0.6698 | 7/0/7 |
| Comp. | RDT-vs-R | NMSE | 0.003 | [0.001, 0.005] | 0.05798 | 4/0/10 |
| Comp. | RDT-vs-RD | PI error (%) | 2.181 | [1.360, 2.997] | 0.001474 | 0/1/13 |
| Comp. | RDT-vs-RD | Flow-rate error (%) | 1.011 | [0.494, 1.580] | 0.004649 | 2/1/11 |
| Comp. | RDT-vs-RD | Peak-velocity error (%) | 0.900 | [0.039, 2.041] | 0.173 | 6/1/7 |
| Comp. | RDT-vs-RD | DivRMS (%) | 0.095 | [-0.210, 0.470] | 0.5067 | 10/1/3 |
| Comp. | RDT-vs-RD | PSNR (dB) | -0.197 | [-0.422, -0.009] | 0.2787 | 6/1/7 |
| Comp. | RDT-vs-RD | SSIM | 0.006 | [0.001, 0.010] | 0.02313 | 10/1/3 |
| Comp. | RDT-vs-RD | NMSE | 0.002 | [0.000, 0.005] | 0.2489 | 6/1/7 |
| Comp. | RDT-vs-RT | PI error (%) | 0.652 | [0.266, 1.199] | 0.002319 | 2/0/12 |
| Comp. | RDT-vs-RT | Flow-rate error (%) | 0.098 | [-0.229, 0.461] | 0.9515 | 7/0/7 |
| Comp. | RDT-vs-RT | Peak-velocity error (%) | 0.486 | [0.116, 0.938] | 0.02454 | 3/0/11 |
| Comp. | RDT-vs-RT | DivRMS (%) | -1.344 | [-1.542, -1.097] | 0.0001221 | 14/0/0 |
| Comp. | RDT-vs-RT | PSNR (dB) | -0.141 | [-0.253, -0.048] | 0.01074 | 4/0/10 |
| Comp. | RDT-vs-RT | SSIM | -0.007 | [-0.011, -0.004] | 0.002319 | 2/0/12 |
| Comp. | RDT-vs-RT | NMSE | 0.002 | [0.001, 0.003] | 0.01343 | 4/0/10 |
| Factorial | 11-vs-10 | PI error (%) | -4.125 | [-5.392, -2.732] | 0.002366 | 12/1/1 |
| Factorial | 11-vs-10 | Flow-rate error (%) | -0.711 | [-1.270, -0.162] | 0.02771 | 10/1/3 |
| Factorial | 11-vs-10 | Peak-velocity error (%) | -1.315 | [-2.506, -0.429] | 0.008775 | 12/1/1 |
| Factorial | 11-vs-10 | DivRMS (%) | -0.402 | [-0.771, -0.101] | 0.0131 | 11/1/2 |
| Factorial | 11-vs-10 | PSNR (dB) | 0.144 | [-0.025, 0.350] | 0.2787 | 8/1/5 |

| Family | Contrast | Metric | Mean diff. | 95% CI | $p$ | W/T/L |
|---|---|---|---|---|---|---|
| Factorial | 11-vs-10 | SSIM | -0.007 | [-0.012, -0.003] | 0.008775 | 1/1/12 |
| Factorial | 11-vs-10 | NMSE | -0.002 | [-0.004, 0.000] | 0.1961 | 8/1/5 |
| Factorial | 01-vs-00 | PI error (%) | -2.235 | [-3.514, -0.849] | 0.006714 | 11/0/3 |
| Factorial | 01-vs-00 | Flow-rate error (%) | 0.094 | [-0.379, 0.556] | 0.7609 | 6/0/8 |
| Factorial | 01-vs-00 | Peak-velocity error (%) | -0.109 | [-0.696, 0.434] | 0.8077 | 6/0/8 |
| Factorial | 01-vs-00 | DivRMS (%) | 0.172 | [0.105, 0.238] | 0.0003662 | 1/0/13 |
| Factorial | 01-vs-00 | PSNR (dB) | -0.058 | [-0.214, 0.095] | 0.5016 | 6/0/8 |
| Factorial | 01-vs-00 | SSIM | -0.009 | [-0.013, -0.004] | 0.004028 | 2/0/12 |
| Factorial | 01-vs-00 | NMSE | 0.001 | [-0.001, 0.002] | 0.5016 | 7/0/7 |
| Factorial | 10-vs-00 | PI error (%) | 2.214 | [1.541, 2.916] | 0.0001221 | 0/0/14 |
| Factorial | 10-vs-00 | Flow-rate error (%) | 0.809 | [0.340, 1.351] | 0.005249 | 3/0/11 |
| Factorial | 10-vs-00 | Peak-velocity error (%) | 0.612 | [-0.106, 1.566] | 0.3258 | 7/0/7 |
| Factorial | 10-vs-00 | DivRMS (%) | -1.774 | [-2.096, -1.402] | 0.0001221 | 14/0/0 |
| Factorial | 10-vs-00 | PSNR (dB) | -0.230 | [-0.412, -0.072] | 0.05798 | 4/0/10 |
| Factorial | 10-vs-00 | SSIM | -0.000 | [-0.003, 0.002] | 0.6698 | 7/0/7 |
| Factorial | 10-vs-00 | NMSE | 0.003 | [0.001, 0.005] | 0.05798 | 4/0/10 |
| Factorial | 11-vs-01 | PI error (%) | 0.325 | [-1.261, 2.002] | 0.9032 | 6/0/8 |
| Factorial | 11-vs-01 | Flow-rate error (%) | 0.003 | [-0.439, 0.456] | 1 | 8/0/6 |
| Factorial | 11-vs-01 | Peak-velocity error (%) | -0.594 | [-1.070, -0.017] | 0.04187 | 10/0/4 |
| Factorial | 11-vs-01 | DivRMS (%) | -2.348 | [-2.571, -2.058] | 0.0001221 | 14/0/0 |
| Factorial | 11-vs-01 | PSNR (dB) | -0.028 | [-0.133, 0.065] | 0.9032 | 8/0/6 |
| Factorial | 11-vs-01 | SSIM | 0.002 | [-0.005, 0.008] | 0.7609 | 7/0/7 |
| Factorial | 11-vs-01 | NMSE | 0.000 | [-0.001, 0.001] | 0.9515 | 8/0/6 |

#### Supplementary Table S12: Fixed-Test-Case PI Results

Supplementary Table S12: Factorial PI error by fixed test case. Values are mean ± SD across 14 seeds; no seed×case pooling was used for inference.

| Case | CFD case | 00 | 01 | 10 | 11 |
|---|---|---|---|---|---|
| C1 | FFF-1 | 22.97 ± 1.19 | 20.40 ± 2.50 | 24.95 ± 0.87 | 21.31 ± 2.54 |
| C2 | FFF | 15.48 ± 1.17 | 13.35 ± 2.43 | 17.57 ± 0.96 | 13.61 ± 2.66 |
| C3 | FFF-9 | 14.50 ± 3.22 | 10.72 ± 4.92 | 17.41 ± 3.36 | 9.80 ± 4.87 |
| C4 | FFF-19 | 11.74 ± 1.29 | 9.34 ± 2.69 | 14.63 ± 0.82 | 10.52 ± 2.64 |
| C5 | FFF-20 | 20.35 ± 1.02 | 18.31 ± 2.09 | 22.19 ± 0.67 | 19.42 ± 1.73 |
| C6 | FFF-29 | 5.22 ± 1.31 | 4.75 ± 1.31 | 6.81 ± 1.50 | 4.14 ± 1.72 |

#### Supplementary Table S13: Reference PI Values

Supplementary Table S13: Conventional nonnegative reference PI values for the six fixed test cases. Plane values are magnitudes of the consistently oriented reference waveforms.

| Case | CFD case | Inlet | Mid | Outlet 1 | Outlet 2 | Plane mean |
|---|---|---|---|---|---|---|
| C1 | FFF-1 | 0.929 | 0.788 | 0.902 | 0.771 | 0.847 |
| C2 | FFF | 0.877 | 0.934 | 0.736 | 1.247 | 0.949 |
| C3 | FFF-9 | 1.006 | 1.018 | 1.246 | 0.863 | 1.033 |
| C4 | FFF-19 | 0.867 | 0.965 | 0.740 | 1.279 | 0.963 |
| C5 | FFF-20 | 0.914 | 0.814 | 0.898 | 0.790 | 0.854 |
| C6 | FFF-29 | 0.841 | 0.808 | 0.747 | 0.890 | 0.821 |

#### Supplementary Note S14: Scheduler, Checkpoint, and PI Sign-Parity Details

**Executed scheduler.** The additional controlled experiments used the same progressive reconstruction schedule in every arm. Actual executed phases were epochs 1–41 (foundation), 42–51 (linear interpolation), 52–141 (physics phase), 142–151 (linear interpolation), and 152 onward (final phase). This one-epoch offset from the nominal configuration milestones arose from scheduler indexing and was common to all compared arms. Spatial Huber regularity was zero throughout the component and factorial experiments.

**Run reuse and historical comparability.** Factorial 00 reused component R and factorial 10 reused component RDT, with identical run/output/checkpoint identities for every seed. Thus, the component and factorial analyses comprise 84 unique trained models rather than 112, and RDT−R and 10−00 are one paired contrast used in two analytical roles. Historical CR and FBA configurations used spatial Huber regularity of 0.1, whereas all factorial conditions used zero to isolate the two factors; factorial 10/11 are not expected to reproduce the historical CR/FBA means. The historical run directories did not archive an executed commit hash, so implementation-version identity between historical and factorial runs cannot be independently verified.

**Seed 047 checkpoint.** Factorial arms 10 and 11 for seed 047 both selected epoch 35, before biomarker activation. Their exact metric tie is therefore expected under the common checkpoint-selection procedure. The seed was retained; no later checkpoint was substituted and no post hoc exclusion was made.

**Historical FBA/CR selected-checkpoint audit.** The selected epochs for the 14 historical seed pairs were: FBA = {145, 220, 175, 180, 35, 145, 185, 180, 145, 45, 160, 160, 150, 195} and CR = {70, 100, 25, 20, 35, 125, 100, 115, 85, 45, 95, 80, 115, 110}, in seed order {021, 031, 037, 041, 047, 053, 059, 061, 067, 071, 079, 083, 089, 099}. One FBA checkpoint (seed 047, epoch 35) was selected before biomarker activation. At epoch 35, all biomarker weights were zero, so the FBA and CR active objectives were identical under the same seed and data order; their archived validation value, test PI (18.327%), and DivRMS (6.303%) were exactly equal. The primary-comparison tie is therefore structural, although bitwise parameter identity is not claimed because retained checkpoint tensors were unavailable. Seed 071 selected epoch 45 in both arms, during the interpolation window when flow and PI supervision remained zero and percentile-velocity supervision had only begun to ramp. Its validation value, PI, and DivRMS were near-identical but not equal, so no exact-tie mechanism is claimed. Win/tie/loss counts define a tie only when the stored numerical paired difference is exactly zero.

**Odd-dimension RCAN target-size fallback.** The RCAN implementation supplied the HR tensor dimensions as `target_size` during training and evaluation and applied nearest-neighbor resizing whenever integer 2× pixel-shuffle output did not exactly match that target. Retained diagnostic arrays verify activation in FFF-9 (LR $16 \times 13 \times 8$ to target $32 \times 27 \times 16$, one-voxel Y correction), FFF-20 ($19 \times 17 \times 11$ to $38 \times 35 \times 22$, one-voxel Y correction), and FFF-29 ($14 \times 14 \times 8$ to $29 \times 28 \times 16$, one-voxel X correction). These cases belong to the fixed test set used by the historical and additional controlled RCAN analyses, confirming that the same evaluation path was exercised across compared arms. The frozen reproducibility archive did not retain spatial-shape metadata for all 22 simulations, so complete prevalence is not claimed. The nearest-neighbor fallback implementation predates the first retained historical primary run.

**PI convention and parity audit.** The manuscript reports conventional nonnegative $PI = (V_{sys} - V_{dia})/|V_{mean}|$. The executed training surrogate and historical PI-validation quantity retained the plane-orientation sign in $V_{mean}$. For the additional controlled analyses, we recomputed conventional and signed PI errors from the underlying plane time curves for 84 unique checkpoints, comprising 2,016 unique run/case/plane observations (2,688 family-labeled observations after accounting for intentional factorial reuse). Prediction and reference mean-flow signs agreed in every observation. The maximum absolute plane-level PI percentage-error difference was $2.1239 \times 10^{-6}$ pp and the maximum arm-mean difference was $1.4164 \times 10^{-7}$ pp. No reported arm mean, paired difference, win/tie/loss count, Wilcoxon statistic or $p$ value, bootstrap interval at reported precision, or interaction conclusion changed. The PI training loss is conditionally invariant when prediction and reference retain the same orientation sign; it is not claimed to be universally sign-invariant. The older dedicated PI-validation reruns remain described according to the orientation-signed criterion actually used, and no counterfactual checkpoint identity is asserted.

**Interpretive scope.** All additional statistics summarize training stochasticity on the same fixed six-case test set. The seed×case observations are retained descriptively and are not treated as independent anatomical samples.

#### Supplementary Figure S1: Historical Exploratory PSNR–PI Scatter

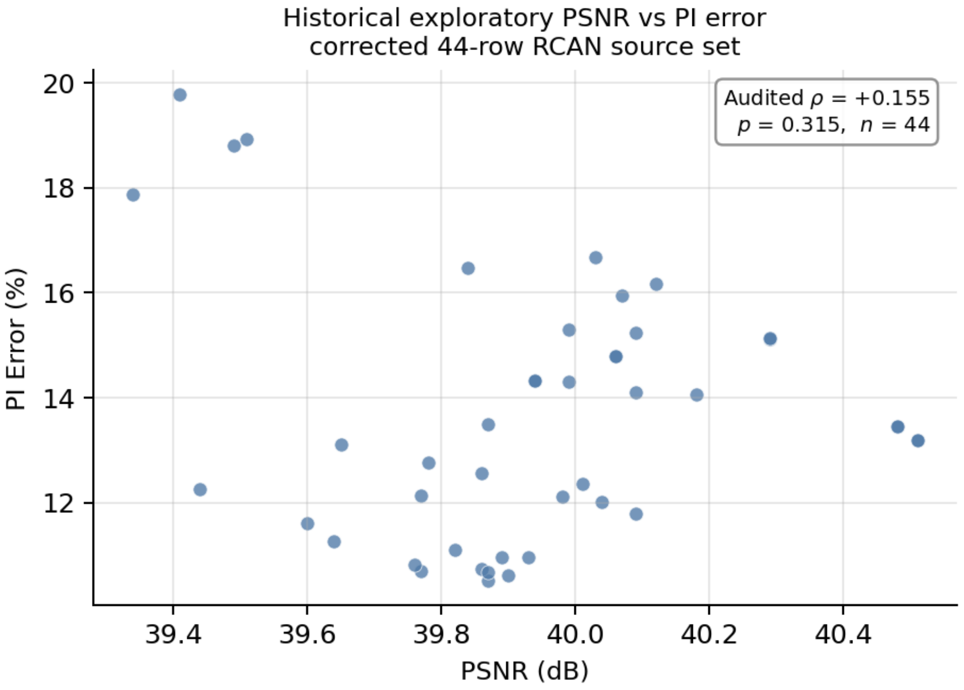


Supplementary Figure S1: PSNR and PI error for the corrected 44-row historical exploratory source set ($\rho = +0.155$, two-sided $p = 0.315$, $n = 44$). Each point is one audited observation in Supplementary Table S7. This figure is retained for provenance transparency and is not evidentiary support for the conclusions.

#### Supplementary Figure S2: Three-Level Ablation — Per-Seed PI and DivRMS Trajectories

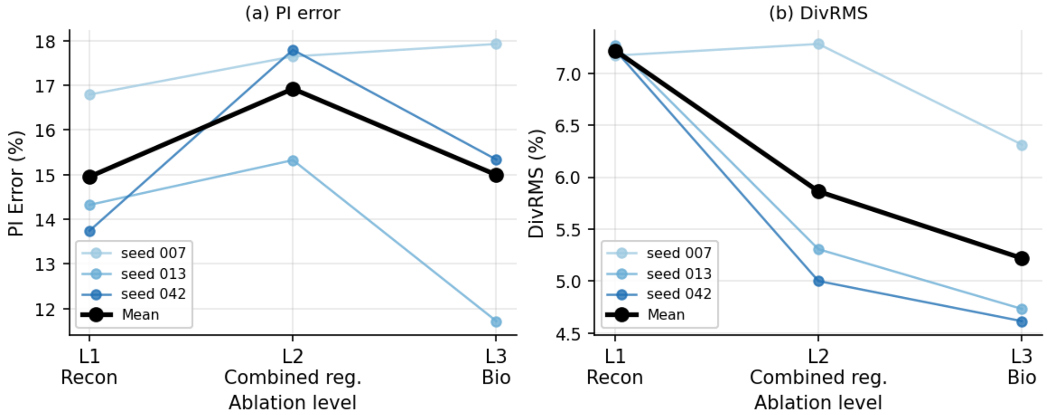


Supplementary Figure S2: Per-seed trajectories and cohort means for PI (left) and DivRMS (right) across the three ablation levels (seeds 007, 013, and 042). DivRMS decreases monotonically at the cohort level (7.22% → 5.86% → 5.22%), whereas the PI response remains heterogeneous: seed 013 shows the full recovery while seeds 007 and 042 do not. Accordingly, within this 3-seed ablation, the evidence is directional for divergence improvement but not for a PI benefit; the primary PI evidence is provided by the 14-pair analysis (main text Section 3.1).

#### Supplementary Figure S3: Paired DivRMS Scatter — Full Biomarker-Aware vs. Reconstruction-Only

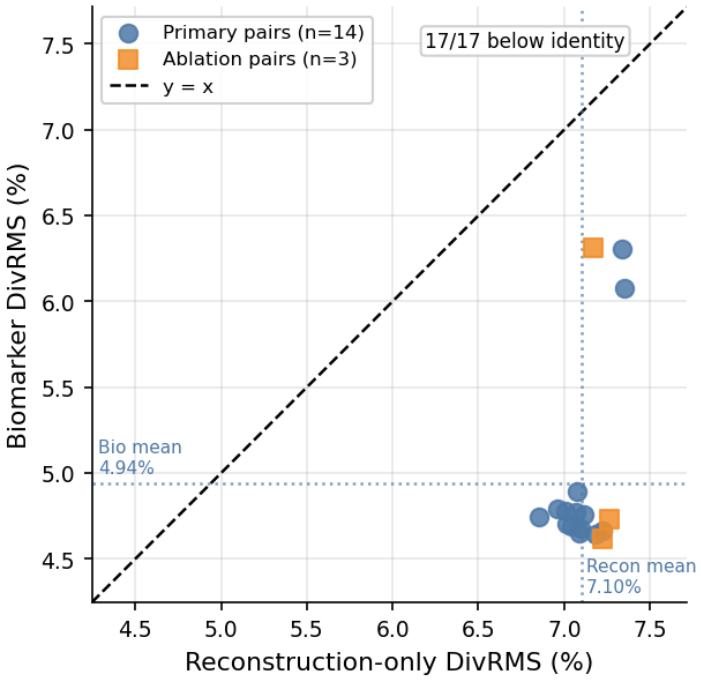


Supplementary Figure S3: Paired DivRMS scatter for all 17 verified full-biomarker-aware vs. reconstruction-only comparisons (14 primary pairs plus 3 ablation Level 3 vs. Level 1 pairs). All 17 points fall below the identity line, indicating lower divergence RMS for the full biomarker-aware arm in every direct comparison within this benchmark. Across all 17 points, mean DivRMS was 4.99% for full biomarker-aware and 7.12% for reconstruction-only; the primary 14-pair cohort means were 4.94% and 7.10%, respectively.
